# LCI9 is required for normal pyrenoid starch sheath formation and efficiency of the CO_2_-concentrating mechanism in *Chlamydomonas reinhardtii*

**DOI:** 10.1101/2025.11.18.689020

**Authors:** Liat Adler, Justin Lau, Lara Esch, Ousmane Dao, James Barrett, Seb Grant, Gary Yates, Alistair J McCormick, Luke CM Mackinder

## Abstract

- Pyrenoid -based CO_2_-concentrating mechanisms (pCCMs) boost photosynthesis by delivering elevated levels of CO_2_ to Rubisco. Pyrenoids are often surrounded by a starch sheath thought to enhance pCCM efficiency, but little is known about how the sheath is formed. Here we have assessed the role of the pyrenoid-associate protein LCI9 in starch sheath formation in *Chlamydomonas reinhardtii*.
- Using an *lci9* mutant we assessed the role of LCI9 in starch sheath morphology by fluorescence and electron microscopy, and its impact on pCCM function by CO_2_ uptake and growth assays. We determined potential interactors of LCI9 via TurboID proximity labeling.
- In the absence of LCI9, cells fail to correctly assemble the starch sheath, have lower affinity for inorganic carbon and grow slower under CO_2_-limiting conditions. LCI9 localises to the starch sheath and is physically close to a range of starch metabolism enzymes and other potential scaffolding proteins. Finally, we show that LCI9 associates with starch in the land plant *Arabidopsis thaliana*.
- LCI9 likely acts as a pyrenoid specific scaffold for starch metabolism enzymes. LCI9 could contribute to ongoing efforts to build a functional pyrenoid in a land plant.

## Introduction

Ribulose-1,5-bisphosphate carboxylase/oxygenase (Rubisco) is the primary carboxylase enzyme for CO_2_ fixation in photosynthetic organisms. Despite its crucial role, Rubisco has a secondary oxygenase activity that limits photosynthetic productivity. To overcome the inefficiencies of Rubisco many photosynthetic organisms have evolved specialized microcompartments called pyrenoids. Pyrenoids are biomolecular condensates consisting of a dense matrix of phase-separated Rubisco. They form the heart of a biophysical CO_2_-concentrating mechanism (CCM) that delivers high concentrations of CO_2_ to Rubisco within the matrix to improve the efficiency of CO_2_ assimilation and suppress oxygenation. Pyrenoid-based CCMs (pCCMs) can elevate the concentration of CO_2_ within the Rubisco matrix to 40-fold above ambient air levels (Badger *et al*.,1980; Fei *et al*., 2022) . Pyrenoids are found in nearly all eukaryotic phytoplankton and are believed to mediate approximately one third of global carbon fixation (Mackinder *et al*., 2016).

In the model green alga *Chlamydomonas reinhardtii* (hereafter Chlamydomonas), the pyrenoid Rubisco matrix is traversed by specialised thylakoid membranes known as pyrenoid tubules that deliver inorganic carbon (Ci) as CO_2_ to Rubisco, and surrounded by a layer of starch known as the starch sheath. Models predict that the starch sheath prevents leakage of CO₂ out of the pyrenoid (Ramazanov *et al*., 1994; Fei *et al*., 2022). Starch sheaths are common in green algal pyrenoids, although their morphology can vary from globular granules loosely associated with the Rubisco matrix, to tightly associated curved plates of starch (Barrett *et al*., 2021). In Chlamydomonas, the starch sheath consists of multiple curved starch plates, known as pyrenoid starch, that surround the pyrenoid with gaps where the pyrenoid tubules access to the matrix. The starch sheath is essential for efficient operation of the pCCM, particularly under very low CO_2_ conditions (<0.01% CO_2_) (Toyokawa *et al*., 2020).

Chlamydomonas also produces starch in the stroma, which is not associated with the pyrenoid. Stromal starch granules are lenticular in shape, similar to those in plant chloroplasts. Under high CO_2_ conditions most starch is located in the stroma in lenticular form, but under ambient or low CO_2_ conditions, starch is mobilised around the pyrenoid as concave plates. Despite these differences in morphology and location, the ratio of amylose and amylopectin in stromal and pyrenoid starch appears to be similar (Izumo *et al*., 2011), although it has been reported that pyrenoid starch may contain slightly more amylopectin (Kuchitsu *et al*., 1988). The formation of stromal starch has been reasonably well studied in Chlamydomonas (Delrue *et al*., 1992; Libessart *et al*., 1995; Courseaux *et al*., 2023; 2025). Comparatively, the synthesis and turnover of pyrenoid starch is less well understood. It is thought that stromal starch could relocalise to the pyrenoid and then be reshaped, and/or that new starch granules could be synthesised and shaped at the site of the pyrenoid. However, the mechanisms by which pyrenoid starch is recruited, shaped into plates and arranged as a sheath are not known.

Recruitment of starch to the pyrenoid appears to be mediated by putative scaffolding proteins SAGA1 and SAGA2 (StArch Granule Abnormal) that have been implicated in mediating adherence of the starch sheath to the Rubisco matrix (Itakura *et al*., 2019; Meyer *et al*., 2020). Both proteins are large coiled-coiled proteins bearing carbohydrate binding motifs (CBM20) and motifs that bind to the small subunit of Rubisco (Mackinder *et al*., 2016, Atkinson *et al*., 2019, He *et al*., 2020, Meyer *et al*., 2020). Loss of SAGA1 appears to cause a broad range of phenotypes. For example, the *saga1* mutant has fewer, thinner and more elongated starch plates and a high CO_2_ requiring phenotype (Itakura *et al*., 2019). Furthermore, *saga1* is characterised by multiple pyrenoids that lack traversing pyrenoid tubules and impaired relocalisation of proteins to the pyrenoid under very low CO_2_ conditions (Shimamura *et al*., 2023). SAGA2 also localises to the starch sheath-Rubisco matrix interface (Meyer *et al*., 2020). More recent data has shown that SAGA1 and SAGA2 are able to recruit starch granules to a ‘proto-pyrenoid’ Rubisco condensate expressed in the model plant *Arabidopsis thaliana* (Arabidopsis) (Atkinson *et al*., 2024).

Apart from SAGA1 and SAGA2, several other proteins associated with pyrenoid starch synthesis have been identified by various approaches, including fluorescent protein tagging (Mackinder *et al*. 2017, Wang *et al*. 2023), pyrenoid protein-protein interaction studies (Mackinder *et al*. 2017) and pyrenoid proximity labeling (Lau *et al*., 2023). Most have yet to be functionally characterised. One such protein is LCI9 (low CO₂-induced protein 9), which was previously identified as a low CO₂ induced gene (Jungnick *et al*., 2014). LCI9 was subsequently confirmed to be a pyrenoid protein that localises to the interface between the plates of the starch sheath (Mackinder *et al*. 2017).

In this study, we found that LCI9 plays a critical role in starch sheath formation. LCI9 is required for normal morphology of the starch sheath and efficient operation of the pCCM. We identified potential interaction partners for LCI9 through TurboID, which include starch synthesis enzymes and putative scaffolding proteins. When expressed in Arabidopsis, LCI9 localizes to the chloroplast and associates with starch granules. Collectively, the data supports LCI9 as an important player in starch sheath formation and a potential target for starch sheath engineering.

## Methods and Materials

### Alphafold structure prediction

Full-length sequences of LCI9 (Cre09.g394473) were submitted to Alphafold Multimer v2 with default settings. The top-ranked model of five was used for figure generation. Figures were prepared using ChimeraX (Pettersen *et al*., 2021).

### Phylogenetic analysis

The evolutionary history was inferred by using the Maximum Likelihood method and JTT matrix-based model (Jones *et al*., 1992). The tree with the highest log likelihood (-45,002.51) is shown. The percentage of trees in which the associated taxa clustered together is shown next to the branches. Initial tree(s) for the heuristic search were obtained automatically by applying Neighbor-Join and BioNJ algorithms to a matrix of pairwise distances estimated using the JTT model, and then selecting the topology with superior log likelihood value. The tree is drawn to scale, with branch lengths measured in the number of substitutions per site (next to the branches). This analysis involved 18 amino acid sequences. There were a total of 3,792 positions in the final dataset. Evolutionary analyses were conducted in MEGA11 (Tamura *et al*., 2021). Chlamydomonas gene sequences were taken from Chlamydomonas genome v5.6.

### Generation of plasmids

The plasmids for fluorescent protein expression and complementation in Chlamydomonas were prepared using a recombineering method described previously by Emrich-Mills *et al*. (2021). For TurboID proximity labelling, the LCI9 (Cre09.g394473) gene sequence is amplified from genomic DNA and cloned into L2-TurboID with PSAD promoter-terminator pair, carrying a paromomycin resistance cassette; L2-RPE1-TurboID (Cre12.g511900) was cloned previously in the same manner as described by Lau *et al*., (2023). For plant-expression, codon optimised and GoldenGate cloning domesticated CreLCI9 (Cre09.g394473) was obtained as a gBlock gene fragment from IDT DNA (Integrated DNA Technologies) and cloned into the level 0 acceptor vector pAGM1287 of the Plant MoClo system (Engler *et al*., 2014). Gene expression constructs of the mCherry tagged protein were assembled into a binary level 2 acceptor. Expression of CreLCI9-mCherry was driven with the Arabidopsis Ubiquitin10 promoter (UBQ10). Plasmids and primers are listed in **Table S1 and S2**.

### Generation of Chlamydomonas Strains

Strains used and generated for this study are listed in **Table S3**. Insertion mutants for the LCI9 locus were obtained from the CLiP library (Li *et al*., 2019). Strains LMJ.RY0402.123483 (*lci9-1*) and LMJ.RY0402.172898 (*lci9-2*). Strains were validated by PCR using primers listed in **Table S2** and sequencing (Fig. **S1**). Transgenic lines were generated by electroporation as described by Yamano *et al*. (2019). Cells were plated for selection on TAP agar supplemented with paromomycin (20 μg mL^−1^) and/or hygromycin (25 μg mL^−1^) under 20 μmol photons m⁻² s⁻¹. Mutants and complemented strains were validated by immunoblot (Fig. **S2**).

### Immunoblot analysis of Chlamydomonas

For immunoblotting, cells were grown photoautotrophically to mid-log phase and were harvested by centrifugation 17,900 × g for 5 min at 4 °C. Cell pellets were resuspended in lysis buffer (25 mM Tris–HCl pH 7.4, 300 mM NaCl, 1 mM DTT, 5 mM MgCl2, 0.1 mM PMSF, 1× EDTA-free protease inhibitor (Roche), 0.1% (w/v) SDS, 0.5% (w/v) deoxycholic acid, and 1% (v/v) Triton X-100) before snap-freezing in liquid nitrogen. The cell suspensions were lysed by 5 freeze/thaw cycles and centrifuged at 17,900 × g for 10 min at 4 °C. The resulting supernatants were used as protein samples in later experiments and stored at −70 °C if not used immediately. For immunoblotting, boiled protein samples were resolved by SDS–PAGE and transferred to a PVDF membrane via a semidry transfer system. The membrane was blocked with 3% (w/v) BSA in Tris-buffered saline with 0.1% (v/v) Tween 20 (TBST) and probed with antibodies accordingly. Antibodies were diluted in TBST as follows: Streptavidin Dylight-488 conjugate (1:4,000, Fisher Scientific #21832); anti-HA (1:1,000, Fisher Scientific 26183); anti-Flag (1:1,000, Sigma #F1804); anti-LCI9 (1:1,000), generated for this study raised to the LCI9 peptide: GDYDLSRRRVDTRVREMQLAK by Yenzym Antibodies; and anti-tubulin (1:2,000, Sigma #T6074).

### Ci Affinity measurements

Ci affinity was estimated from O₂ evolution rates at varying Ci concentrations using a Clark-type oxygen electrode (Oxygraph Plus, Hansatech). Autotrophic cultures grown at 3% or 0.04% CO₂ were harvested by centrifugation (1,000 × g, 5 min, 25 °C) and resuspended to 20 μg Chl mL⁻¹ in 2 mL CO₂-depleted TP medium. Measurements were performed in the electrode chamber equilibrated to 25 °C, stirred at 75%, and illuminated at 500 μmol photons m⁻² s⁻¹. Cultures were bubbled with CO₂-free air until the net O₂ evolution rate was ∼0. NaHCO₃ was sequentially added (1–500 μM) using a Hamilton syringe, and O₂ evolution was recorded for 1 min after each addition. Rates were calculated over a 40 s window.

### Chlamydomonas Growth Assays

Spot Tests: Cells were grown mixotrophically in TAP media. Once cultures reached 2−4 × 10⁶ cells mL^−1^, 1 × 10⁶ cells were harvested by centrifugation at 1,000 x g for 10 min. Cells were washed and resuspended at a concentration of 1 × 10⁶ cells ml^−1^ in TP media. Cell concentration was measured using Countess® II FL Automated Cell Counter (Thermofisher). Liquid cultures were spotted onto TP agar (1.5%) in 1,000, 100 and 10 cell spots at a range of pHs (specified). The plates were incubated in 3%, 0.04% and 0.01% CO₂ and illuminated under constant light at 400 μmol photons m^−2^ s^−1^. Growth was monitored for up to 10 days and photos were taken using the PhenoBooth+ Colony counter (Singer).

### Confocal Microscopy

Transgenic fluorescent strains were initially grown mixotrophically in TAP media until reaching 2 –4 ×10^6^ cells mL^−1^ and resuspended in TP media overnight prior to imaging. Cells were mounted on 8-well chamber slides and overlaid with 1.5% (w/v) low melting point agarose made with TP-medium. Images were collected on a LSM880 or LSM980 (Zeiss) equipped with an Airyscan module using a 63× objective. Laser excitation and Emission setting for each channel used are set as below: Venus(excitation: 514 nm; emission 525 – 500 nm); mScarlet-I (excitation: 561 nm; emission 570 – 620 nm); Chlorophyll (excitation: 633 nm; emission 670 – 700 nm). Analysis of starch plate morphology was performed using the freehand selection tool and ‘Measure’ function in Fiji (Schindelin *et al*., 2012).

### Streptavidin-affinity purification

Biotinylated proteins were enriched via streptavidin-affinity purification as previously mentioned (Lau *et al*., 2023) with minor modifications. Wildtype (cc-5325) cells and cells expressing RPE1-TurboID or LCI9-TurboID were inoculated into 400 mL TP-medium in a Duran bottle bubbled with 3% CO_2_ supply and with constant stirring at 160 rpm. Cells were allowed to grow until 0.5 – 1×10^6^ cells mL^−1^ before being switched to 0.04% CO_2_ supply for at least 2 days to induce the CCM. After CCM induction, cells are harvested by centrifugation and resuspended in TP-medium to an OD750 of 2.5. The suspensions were then split into triplicates before biotin (in DMSO) was added to the cell culture to a final concentration of 2.5 mM. Biotin incubation step was allowed to proceed for two hours and halted by centrifugation 21,300 × g, 2 min at 4 °C. Biotin-labelled cells were rinsed three times with ice-cold TP-medium before being snap-frozen in liquid nitrogen until lysis. Cell lysis was performed as mentioned above except protein loading dye was not added to the clarified cell lysate. Zeba Spin desalting columns (#89891, Thermo Fisher) were used to remove any free biotin in lysate. The BCA protein assay (ThermoFisher) was used to measure protein concentrations after diluting protein extract 1:10 and 1.75 mg protein was added to the 50 μL Streptavidin beads that was pre-equilibrated with lysis buffer. Overnight incubation of beads with protein lysate was carried out in a 4°C cold room with constant rotation. Beads were then washed twice with lysis buffer for five minutes: once with 1 M KCl for 2 min; once with 0.1 M NaCO_3_ for 1 min; once with 4 M urea in 50 mM triethylammonium bicarbonate, pH 8.5 (TEAB) for 1 min; once with 6 M urea in 50 mM TEAB for 1 min; and twice with 50 mM TEAB buffer for 5 min. The washed beads were frozen at −70 °C until they were submitted for mass spectrometry.

### Mass spectrometry analysis

The TurboID samples were analysed with a TimsTOF HT mass spectrometer (Bruker). On-Bead digestion of peptides was performed as previously described (Lau *et al*., 2023). EvoSep LC separation was performed using the 30 SPD pre-set elution method and a C18 Performance column (8 cm x 150 mm x 1.5 μm). The nanoUPLC was interfaced to a timsTOF HT mass spectrometer (Bruker) with a CaptiveSpray ionisation source (Source). Positive PASEF-DIA (Parallel accumulation serial fragmentation – Data independent acquisition), spectra were acquired using Compass HyStar software (version 6.2, Thermo). Instrument source settings were as follows: capillary voltage, 1,500 V; dry gas, 3 L min^−1^; dry temperature; 180°C. The PASEF mass spectrometry method used a 1.8 s cycle time with MS2 DIA windows of 50 m/z between m/z 400-1201 and a mobility range of 0.6-1.43 1/k0. The fragment scan m/z range was 100-1700. Resulting data were searched using DIA-NN (1.8.2.27) against the Chlamydomonas sequences from the *Chlamydomonas reinhardtii* genome cc-4532 v6.1 (Joint Genome Institute, https://jgi.doe.gov/) appended with common proteomic contaminants and the TurboID protein. The search was performed at 1% false discovery rate with data filtered to require individual protein q-values as less than 0.01 for identification and a minimum of two peptides per accepted protein. DIA-quantification was performed within DIA-NN. Statistical comparison between sample groups was performed using an in-house installation of FragPipe-Analyst within R Shiny. Using FragPipe-Analyst pairwise comparisons were conducted between all groups with sample minimum imputation applied. Non-adjusted p-values are used to plot volcano plots.

### Starch granule purification, morphology and size analysis for Chlamydomonas

Cells were adapted to 0.04% CO_2_ for at least 48 hours before being harvested. Starch was isolated using an adapted protocol from Delrue et al. (1992). Briefly, cells were collected by centrifugation at 1000 x g for 5 min and resuspended in starch extraction buffer (SEB) (50 mM Tris-HCl (pH 8), 0.2 mM EDTA and 0.5% (v/v) Triton X-100) at 10^8^ cells mL^−1^. Cells were sonicated on ice for 3 min at amplitude 30, 3 s on 10 s off using a Misonix S-4000. Five mL of crude extract was placed on a 5 mL percoll cushion (95% (v/v) Percoll/1x SEB) and centrifuged at 2,500 x g for 15 min at 4 ℃. The white pellet containing isolated starch was resuspended in 5 mL dH_2_O and passed through the percoll cushion again. The final white pellet containing starch was weighed and resuspended at 10 mg mL^−1^ and 4 mg mL^−1^ in dH_2_O for further analysis by scanning electron microscopy (SEM) and particle analyzer, respectively. For SEM, the samples of the starch suspensions were vortexed and 9 µL aliquots were dried down onto pieces of silica. These were then affixed to SEM stubs and sputter coated with 5 nm of gold/palladium on a Polaron SC7640 sputter coater. Then imaged using Jeol JSM 6490LV scanning electron microscope operating at 10 kV accelerating voltage and WD of 15 mm. Representative images were collected at 2K, 5K and 10-20K as applicable. For particle size analysis, starch was resuspended at 5 mg mL^−1^ in distilled water for analysis. The size distribution of extracted starch granules was determined using a dynamic light scattering instrument Zetasizer Nano (Malvern).

### Transmission electron microscopy of Chlamydomonas

For low CO_2_ conditions, cells were bubbled with 0.04% CO_2_ gas mixture overnight,harvested via centrifugation (×1,000 g, 4 min, at room temperature), then fixed in 2.5% (v/v) glutaraldehyde in TP-medium for 1 hour at room temperature with constant rotation. Fixed cells were washed for a minimum of 10 min before incubation in 1% (w/v) OsO_4_, 1.5% (w/v) K_3_[Fe(CN)_6_] on ice, in the dark, for 1-2 h. The osmicated samples are washed with dH_2_O and resuspended in low melting point agar in (3% (w/v) in dH_2_O) to give a final agar concentration of 1-2% (w/v). Agar blocks (max 2 mm^2^) containing cells were dehydrated through a graded ethanol series (30%, 50%, 70%, 90% (v/v) for 10 minutes at each concentration, followed by two 15-minute incubations in 100% ethanol). Cells were then infiltrated with Spurr replacement resin (TAAB Laboratories Equipment Ltd, England); resin: ethanol, 1:2. 1:1. 2:1, followed by three incubations in 100% resin. Each resin incubation was a minimum of 1 hour and at least one 100% incubation was overnight. Samples were embedded in flat embedding moulds and resin was polymerised at 70°C for 48 hours. Ultrathin sections (70 nm) were cut using a Leica EM UC7 ultramicrotome and collected on uncoated copper grids (200 mesh). Sections were post-stained with 2% (w/v) uranyl acetate in 50% (v/v) ethanol in the dark, followed by staining with Reynold’s lead citrate (Reynold, 1963) in a CO_2_-depleted chamber, both steps carried out for 5 minutes. Images were acquired using a Jeol JEM-1400 Transmission electron microscope operating at 120 kV (Jeol USA).

### Plant material and growth conditions

Arabidopsis thaliana wildtype (Col-0) and transgenic plants were grown under controlled environment conditions (21° C and 70% relative humidity and 150 µmol m ^−2^ s ^−1^) in 12 hours light 12 hours dark cycles.

### Arabidopsis plant transformation

Plasmids were transformed into *Agrobacterium tumefaciens* (AGL1) for plant transformation. Arabidopsis Col-0 plants were transformed via floral dip (Zhang *et al*., 2006) and three independent transgenic lines were identified using the pFAST-R selection (Shimada *et al*., 2010). Experiments were performed with plants in the T2 generation.

### Confocal microscopy for Arabidopsis

Confocal laser scanning images of WT and UBQ10:CreLCI9-mCherry mesophyll cells were taken in the middle of the photoperiod of 21 day-old plants using a Leica SP8 confocal microscope and a 63x water dipper objective. mCherry fluorescence was excited at 552 nm and emission was detected at 593-620 nm. Chlorophyll fluorescence was excited at 488 nm and emission was detected at 650-700 nm.

Fluorescein staining was performed using a 100 µM solution adapted from Ichikawa *et al*. (2024) and fluorescence was excited at 488 nm and detected at 503-532nm. Images were processed using ImageJ software (http://rsbweb.nih.gov/ij/) and Adobe Photoshop 2024.

### Starch granule purification, morphology and size analysis for Arabidopsis

Four-week-old plants (60 rosettes per genotype) were pooled and homogenised in 50 mM Tris-HCl, pH 8.0, 0.2 mM EDTA, 0.5% (v/v) Triton X-100 using an immersion blender. The suspension was sequentially filtered through Miracloth, a 60 µm nylon mesh and a 20 µm nylon mesh. Starch granules were separated by centrifugation at 2,500 x g over a Percoll cushion (95% (v/v) Percoll, 5% (v/v) 0.5 M Tris-HCl, pH 8.0) then washed in water until clean, then washed twice in 0.5% (w/v) SDS in water and once in 100% ethanol to dry. Starch granule morphology was imaged using a ZEISS Crossbeam 550 SEM. Granule size distribution was analysed by measuring the diameter of 200 starch granules per genotype from SEM images using ImageJ. Images were processed using ImageJ software (http://rsbweb.nih.gov/ij/) and Adobe Photoshop 2024.

### Starch binding assay, protein extraction and immunoblotting for Arabidopsis

To assess starch binding, total protein was extracted from Arabidopsis leaves in 40 mM Tris HCl, pH 6.8, 5 mM MgCl_2_, 1 mM DTT and 1x Proteinase inhibitor Cocktail (PiC). Input samples were taken after centrifugation at max speed at 4°C and protein extracts were incubated with purified starch granules for 1 h at 8°C rolling. Output samples were taken and starch was pelleted and washed 3x in the protein extraction buffer. Then, the starch pellet was incubated for 30 min at 37°C in 50 µl 40 mM Tris, pH 6.8, 5mM MgCl_2_ 2xPiC and 4% (w/v) SDS, while shaking, to elute starch bound proteins for eluted samples. To assess CreLCI9-mCherry expression in transgenic lines, proteins were extracted in 50 mM Tris HCl, pH8, 150 mM NaCl, 1% (w/v) Triton × 100, 1xPiC and 1 mM DTT. For immunoblotting, the anti-mCherry antibody (abcam, ab213511) was used in a 1:2,000 concentration. Bands were detected using chemiluminescence, using the Goat pAb to Rb IgG (HRP) (abcam, ab6721– 1:10,000) and the Pierce™ ECL Western Blotting Substrate (Thermo Fisher Scientific).

## Results

### LCI9 is located to the periphery of pyrenoid starch plates

LCI9 has two predicted CBM20 domains in the N-terminus linked by a disordered domain to a coiled-coiled domain in the C-terminus (Fig. **1a**). Previous work has shown that LCI9 localizes in a mesh-like structure around the pyrenoid, which appears to correlate with the interface between the starch sheath plates (Mackinder *et al*., 2017). To confirm this, we generated a dual-tagged line expressing LCI9-mNeonGreen and the granule-bound starch synthase STA2 (STA2-mCherry) (Cre17.g721500) as a starch marker. With this line we found that LCI9 is localised around the periphery of the starch plates and is enriched in the interface where two plates meet (Fig. **1b**).

**Figure 1.**
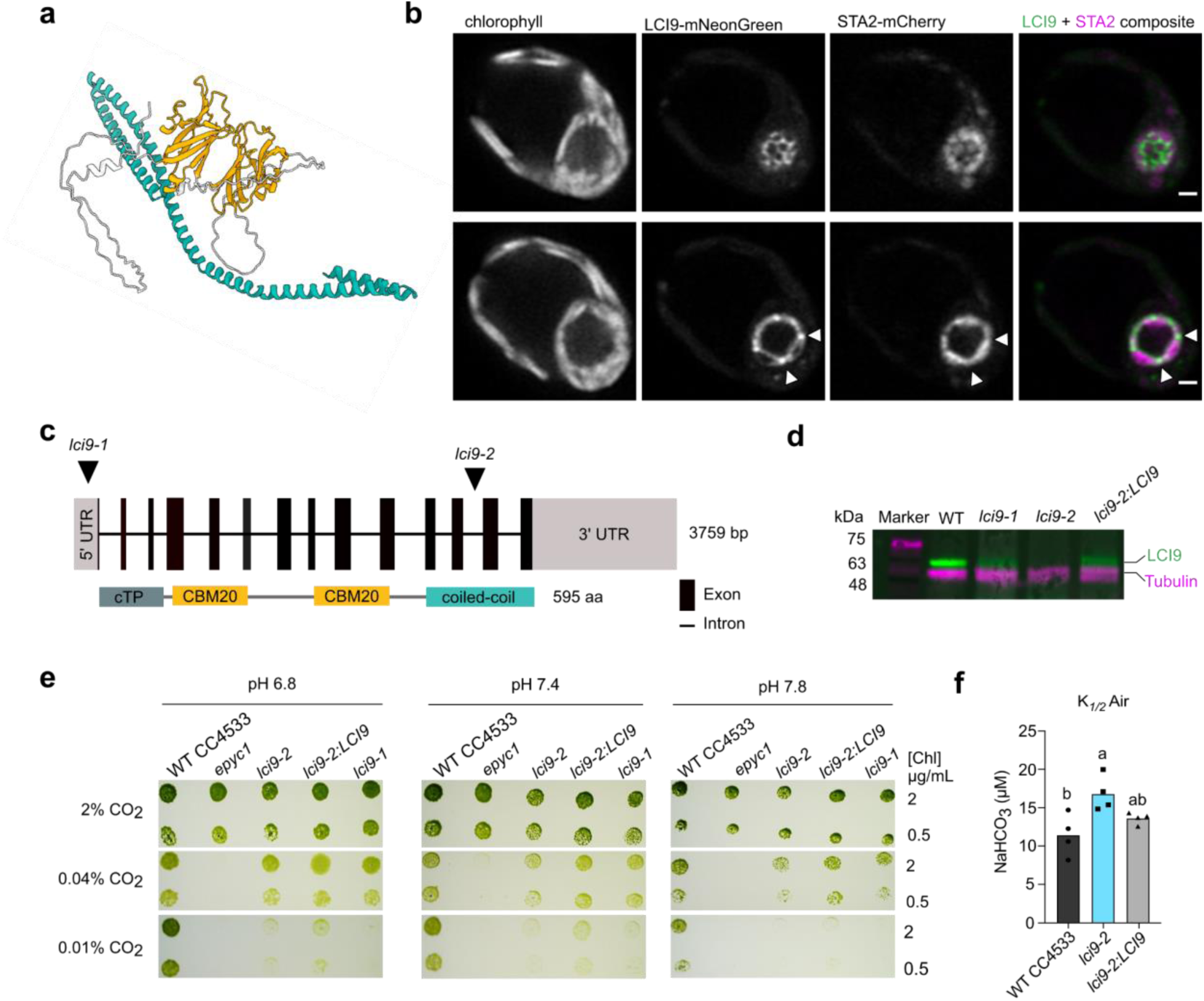
LCI9 localizes to the pyrenoid starch sheath and is required for optimal pCCM operation. **(a)** Alphafold model of LCI9 protein structure. Carbohydrate binding domains (CBM20) are shown in orange and the coiled-coiled region in teal. **(b)** Confocal micrograph of a Chlamydomonas transformant expressing fluorescently tagged STA2-mCherry and LCI9-mNeonGreen shown in magenta and green, respectively. Two z planes of the same cell are shown, scale bar is 2 µm. Taken one hour after transition from high to low CO_2._ Arrows indicate where LCI9-mNeonGreen signal is enriched between starch plate junctions. **(c)** Schematic of the LCI9 gene and the site of CIB1 cassette (contains a paromomycin resistance construct and a unique internal barcode) insertions (top). Schematic of LCI9 protein with domains indicated including the chloroplast transit peptide (cTP) (bottom) **(d)** Immunoblot assessing the accumulation of LCI9 protein in the insertional mutants *lci9-1* and *lci9-2* and the complemented line *lci9-2:LCI9*. **(e)** Solid media growth assay under the indicated CO_2_ and pH conditions. **(f)** Ci affinity of *lci9-2* compared to control strain and complemented line. Different letters indicate significant differences (p<0.05) as determined by a one-way ANOVA.

### LCI9 insertion mutant has a CCM defect

To understand the role of LCI9 in the CCM, we obtained two CLiP mutants, *lci9-1* and *lci9-2* (Li *et al*., 2019) and confirmed cassette insertion via sequencing (Fig. **S1**)*. lci9-1* has a cassette insertion in the 5’UTR and *lci9-2* has an insertion in the 12th intron, which could destabilise mRNA and/or disrupt correct translation of the C-terminal coiled-coil domain (Fig. **1c**). We found from immunoblotting that *lci9-1* was likely a knock-down line as there was still detectable LCI9 (Fig. **1d**). Conversely, no LCI9 was detected in *lci9-2* (Fig. **1d****; S2**). As the ⍺-LCI9 antibody was raised against the C-terminus of LCI9 it was not sufficient to confirm whether *lci9-2* was a full knockout or expresses a truncated version of LCI9. Since *lci9-1* is a knockdown, we used the *lci9-2* mutant for further investigations. We generated a complemented line by expressing LCI9 under its native promoter in the *lci9-2* background. LCI9 expression was restored to 76% of WT in *lci9-2:LCI9* (Fig**. 1d****; S2**).

We assessed the CCM fitness of the mutants and the complemented line via a spot test growth assay on solid agar media under different CO₂ and pH conditions. WT strain CC-4533 (the CLiP library background strain) was used as control. The mutant *epyc1*, a well-characterised CCM deficient mutant, was used as a negative control. The growth of both *lci9* mutants was moderately impaired under air levels of CO₂ (0.04%) and severely impaired at <0.01% CO₂. The growth defect was also worse at pH 7.8 compared to pH 6.8 and pH 7.4 (Fig. **1e**). At higher pH values, a greater proportion of Ci exists as bicarbonate, making cells more reliant on bicarbonate-specific uptake strategies. In contrast, growth of the *lci9* mutants was similar to WT when supplemented with 2% CO₂. Complementation with *LCI9* in the *lci9-2* background partially restored growth under limiting CO₂ conditions. Measurements of Ci affinity with an O₂ electrode showed that *lci9-2* has a lower Ci affinity during CCM operation, which was also partially rescued by complementation with *LCI9* (Fig. **1f**; Fig. **S3**). In summary, growth and Ci affinity measurements support that LCI9 is required for a fully functional CCM.

### lci9-2 has an abnormal starch sheath

Since LCI9 contains CBM20 domains and localises to the starch sheath, we reasoned that the observed CCM phenotype in *lci9-2* may be due to a defect in the starch sheath, thus we compared the morphology of the starch sheath in WT and *lci9-2*. First, we used fluorescently tagged STA2 (STA2-mCherry) as a starch marker combined with super-resolution confocal microscopy to visualise the starch sheath in live cells under low CO₂. In WT, a z-slice showed the starch sheath was uniformly circular (Fig. **2a**). Conversely, *lci9-2* had a disrupted starch sheath with individual starch plates often overlapping each other or clashing (Fig. **2a,b****, S4**). The starch sheath was also squashed in appearance and had reduced circularity (Fig. **2a,c****, S4**). To gain further detail, we obtained transmission electron microscopy (TEM) micrographs of WT, *lci9-2* and *lci9-2:LCI9*. Similarly to what we observed through confocal microscopy, the starch sheath in WT had a canonical circular shape whereas *lci9-2* had an irregular sheath with multiple plate clashes (where the plates overlap, seen as plates not meeting end to end in 2D) and reduced circularity (Fig. **2d****, S5**). The TEM images also showed that the morphology of the starch sheath in *lci9-2:LCI9* was restored to that of the WT. In WT cells under high CO₂, starch granules associated with the pyrenoid were more globular compared to low CO_2_, as has been previously observed (Fig. **S6**) (Ramazanov *et al*., 1994). Starch granules in *lci9-2* were also more globular under high CO_2_ although granules tended to be more elongated than in WT (Fig. **S6**). Surprisingly, the starch sheath in *lci9-2:LCI9* appeared fully formed under high CO_2_ suggesting transgenic expression of LCI9 may interfere with starch sheath breakdown. Together this data supports that LCI9 is important for shaping the starch sheath under low CO_2_ conditions and may inhibit starch sheath breakdown, consistent with LCI9 being a low CO_2_ induced gene.

**Figure 2.**
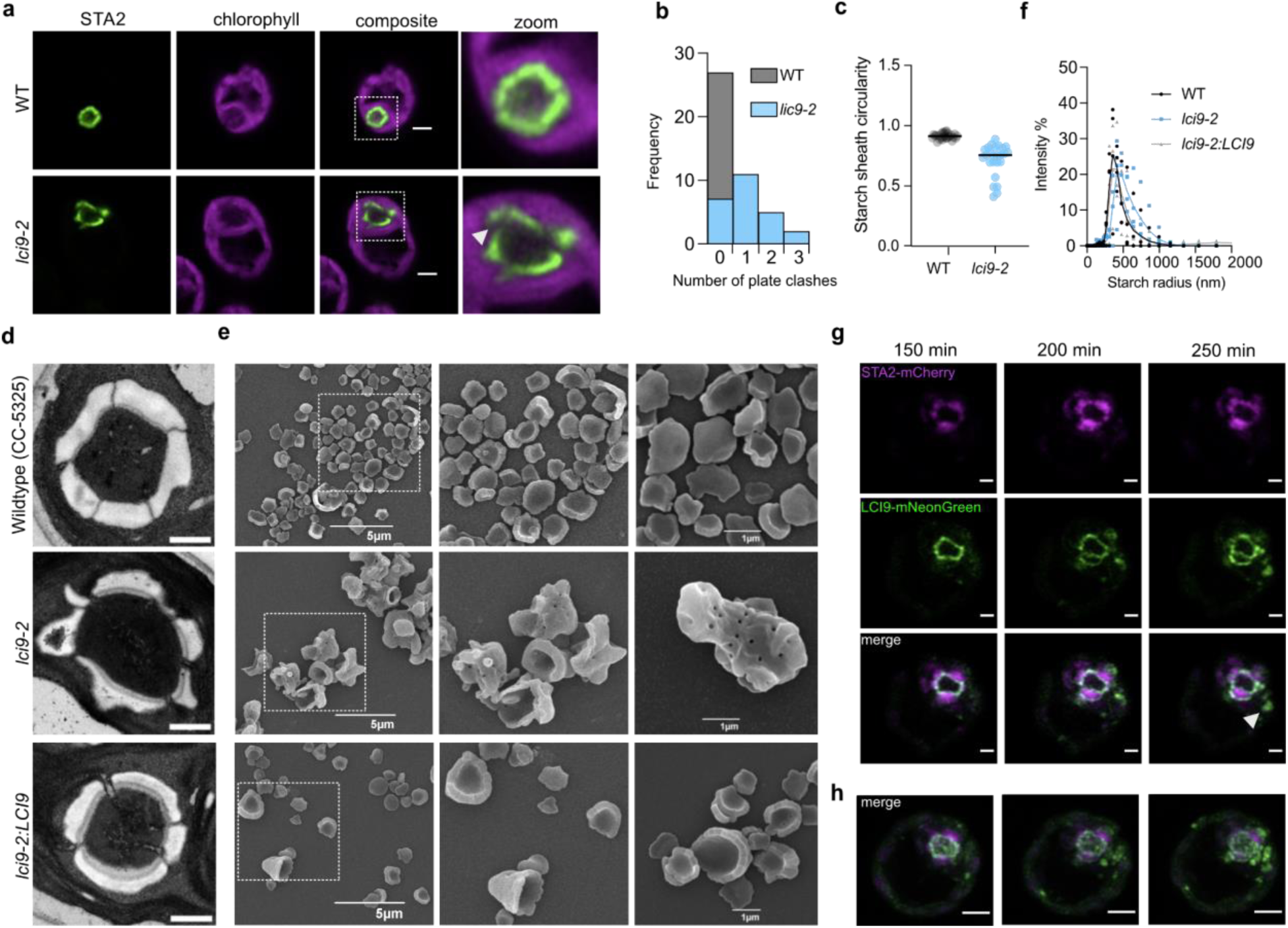
LCI9 is required for normal starch sheath formation. **(a)** Confocal images showing single chlamydomonas cells with STA2-mScarlet-I as a starch marker. **(b)** Number of plate clashes in the starch sheath observed in the middle z-plane of a pyrenoid (n=25). **(c)** Circularity of the starch sheath in WT vs *lic9-2* cells. **(d)** Transmission electron micrographs of pyrenoids from WT, *lci9-2* and complemented strain *lci9-2:LCI9*. Scale bar is 2 µm. **(e)** Scanning electron micrographs of starch extracted from WT, *lci9-2* and complemented strain *lci9-2:LCI9*. **(f)** Quantification of radii of starch particles extracted from low CO_2_ grown cells. Markers represent the mean of three technical replicates for each biological replicate (n=4), line represents the mean of biological replicates. **(g)** Time lapse of cell transitioning from high CO_2_ (3%) to low CO_2_ (0.04%). LCI9-mNeonGreen and STA2-mScarlet-I are shown in green and magenta, respectively. A single z plane through the middle of the pyrenoids is shown. White arrow indicates where the LCI9 signal is observed, seemingly unattached to STA2-labelled starch. Representative of n=8. Scale bar is 1 µm. **(h)** Max z projection of the cell in (g). Scale bar is 2 µm.

To assess the morphology of the individual starch plates, we visualised isolated starch granules by SEM. Starch granules from low CO₂ grown WT cells, when most starch is pyrenoid-associated, were uniform concave discs as previously observed in Izumo *et al*. (2011) (Fig. **2e****; S7**). Conversely, granules from *lci9-2* were non-uniform and varied in shape and size (Fig. **2e****; S7**). The surface of the starch from *lci9-2* was more featured and uneven with some granules having pits or protrusions on the surface. Similarly to WT, the shape of starch granules from the complemented line *lci9-2:LCI9* were uniform concave discs (Fig. **2e****; S7**). The size of starch granules isolated from *lci9-2* were larger than those from the WT control, while the size distribution of granules isolated from *lci9-2:LCI9* was similar to WT (Fig. **2f**). WT and *lci9-2:LCI9* starch granules were also more uniform in size compared to *lci9-2*, which had a wider distribution. The abnormal morphology of starch granules isolated from *lci9-2* suggests that LCI9 has a role in the normal formation of the starch sheath plates.

To gain insight into how LCI9 might influence sheath formation, we performed time-lapse imaging of LCI9-mNeonGreen and STA2-mCherry during the high CO₂ to low CO₂ transition. In our LCI9-mNeonGreen and STA2-mCherry dual-tagged line, LCI9 is under its native promoter that is low CO_2_ regulated (Jungnick *et al*. 2014) whilst STA2 is driven by the constitutive PSAD promoter. In the first hour after switching cells to a low CO₂ environment, LCI9-mNeonGreen expression increased. In several cells, LCI9-mNeonGreen appeared to localise preferentially to the surfaces of starch adjacent to the pyrenoid matrix, before spreading around each starch granule and finally becoming enriched in the interface between the starch plates after approximately four hours (Fig. **2g,h****; S8**). At steady state, LCI9 is localised all around the starch sheath plates and is enriched at the interface between each plate (Mackinder *et al*., 2017).

Interestingly, in some cells we observed hollow puncta of LCI9-mNeonGreen signal adjacent to the pyrenoid later in the timelapse that were not associated with STA2-labelled starch. This may represent LCI9 associating with *de novo*-synthesised starch that is not yet associated with STA2-mCherry activity. The changing distribution of LCI9 during starch sheath formation suggests that LCI9 has a role in shaping the starch around the pyrenoid and may have more of a stabilising role after formation.

### LCI9 may regulate starch enzymes to promote the correct formation of the starch plate sheath

Since LCI9 does not contain any predicted enzymatic domains, we reasoned that LCI9 may influence starch morphology by recruiting starch enzymes. A class of proteins known as Protein Targeting to STarch (PTST) also contain CBMs and coiled-coiled domains like LCI9, and are known to target starch enzymes to starch granules (Seung *et al*. 2015; 2017; 2018; Kamble *et al*., 2023). In a previous study, LCI9 co-immunoprecipitated with starch branching enzyme 3 (SBE3) and two glycolysis enzymes, phosphofructokinase 1 (PFK1) and PFK2 (Mackinder *et al*., 2017). To verify these potential interactors and to identify additional LCI9 associated proteins, we employed a TurboID labeling system developed by Lau *et al*. (2023). TurboID biotinylates proteins in close proximity to a protein of interest, enabling streptavidin purification and protein identification via mass spectrometry to generate a ‘proxiome’. We generated a proxiome for LCI9 under low CO_2_ conditions when the CCM is active and LCI9 is associated with the starch sheath. We used two controls: WT cells where endogenous biotin ligases result in background biotinylation; and TurboID fused to an abundant stromally localised Calvin-Benson-Bassham cycle enzyme ribulose phosphate-3-epimerase (RPE1-TurboID) (Meloni *et al*., 2024)(Fig. **3a,b****; S9**). To generate a high confidence proxiome, we only considered proteins enriched in both LCI9 vs WT and LCI9 vs RPE1, which yielded 24 proteins (Fig. **3c,d**; Table **S4,S5**). In agreement with Mackinder *et al*. (2017), SBE3 was present in the high-confidence LCI9 proxiome (Fig. **3c**). However, PFK1 and PFK2 were not enriched in the LCI9 proxiome and were only enriched in the LCI9/WT comparison group (Table **S4,S5**). To investigate this further, we localised PFK1 and PFK2 using fluorescent tags. Although PFK1 and PFK2 are part of the pyrenoid proteome generated by Zhan e*t al*. (2018), we found that PFK1 and PFK2 do not localise to the starch plates, suggesting that either they may not be true interactors of LCI9 *in vivo,* the interaction is transient, or the fluorescent tag has affected the localisation of the PFKs (Fig. **S10**).

**Figure 3.**
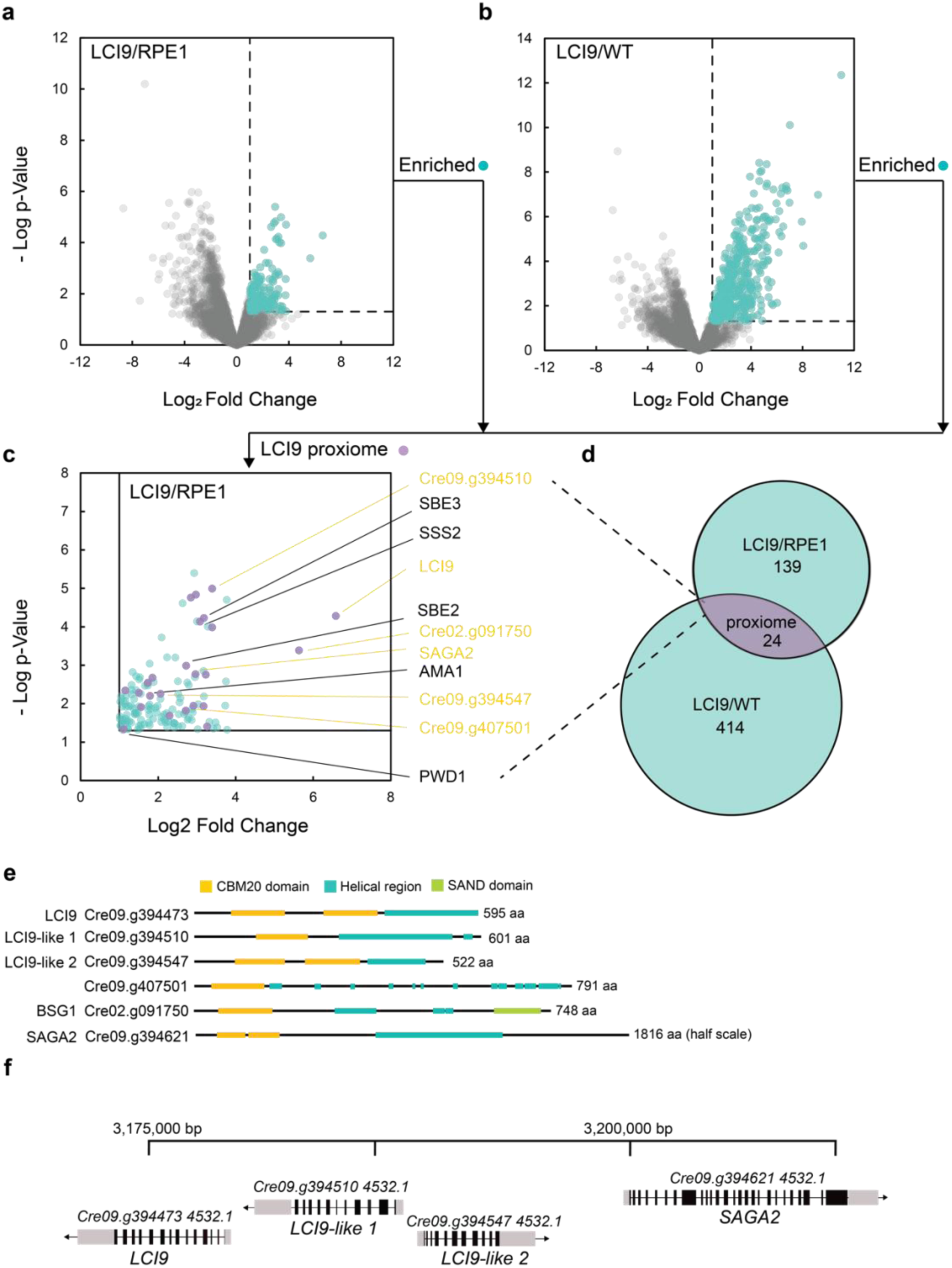
LCI9 proxiome. Preferentially labelled proteins from LCI9-TurboID are compared to RPE1-TurboID **(a)** or wildtype **(b)**. Log2 Fold Change of protein abundance was calculated for detected proteins and visualised as a scatterplot against the Log p-value. The enrichment threshold was set at Log_2_ Fold Change >1 and p <0.05, as shown by the dotted lines. Proteins considered significantly enriched are coloured in teal. **(c)** To obtain a more stringent LCI9 proxiome we found proteins that were enriched in both LCI9/RPE1 and LCI9/WT datasets. These proteins are shown in purple and starch-related proteins are highlighted (starch synthesis/degradation: black; CBM-containing: orange). **(d)** Venn diagram demonstrating the comparison between significantly enriched proteins in each comparison group. **(e)** Annotated domains of each protein of the LCI9 gene cluster are displayed on the schematic of the amino acid sequences on the right. **(f)** Schematic of the LCI9 gene cluster on chromosome nine of the *Chlamydomonas reinhardtii* genome v6.1.

In addition to SBE3, the starch biosynthesis enzymes soluble starch synthase II (SSS2), SBE2, alpha-amylase 1 (AMA1) and phosphoglucan water dikinase 1 (PWD1), were high-confidence components of the LCI9 proxiome (Fig. **3c,d**). We also detected SAGA2, which localises to the starch-matrix interface (Meyer *et al*., 2020), as well as three additional CBM20 domain proteins. One of these, bimodal starch granule 1 (BSG1; Cre02.g091750), has previously been associated with the transition from pyrenoid to stromal starch (Findinier *et al*., 2019). The other two are uncharacterised (Cre09.g394510 and Cre09.g394547), which, together with SAGA2, appear in the same genome cluster as *LCI9*, suggesting that LCI9 might be part of a larger family of related proteins (Fig. **3e,f**). The proteins have between 22-28% amino acid percentage identity with each other (Fig. **S11**). To assess the relationship of this cluster to other CBM-coiled-coiled proteins we performed a phylogenetic analysis (Fig. **S11**). We found that the genes in the *LCI9* gene cluster were more closely related to each other than the Chlamydomonas PTST homolog, suggesting that they have distinct functional roles from PTST. Taken together, LCI9 associates with both starch synthesis and degradation enzymes as well as other starch-related proteins.

### LCI9 localises to the chloroplast and binds to starch in plants

Recently, significant progress has been made toward introducing a functional pCCM into land plants (Atkinson *et al*., 2020; Hennacy *et al*., 2024), including the association of starch granules around a synthetic ‘proto-pyrenoid’ expressed in Arabidopsis (Atkinson *et al*., 2024). Our data shows that LCI9 is necessary for correct starch sheath formation and thus could be a key component to progress pCCM engineering efforts *in planta*. As an initial step, we decided to investigate if LCI9 associates with starch granules in the model C3 plant Arabidopsis. We expressed LCI9-mCherry in WT (AtCol-0) Arabidopsis and visualised the localisation of LCI9 relative to starch by confocal microscopy using fluorescein to stain starch (Fig. **4a,b**; **S12**) (Ichikawa *et al*., 2024). LCI9-mCherry localised clearly around starch granules in mesophyll cell chloroplasts (Fig. **4a,b**). To confirm LCI9 was binding to starch, total protein extracts from LCI9-mCherry expressing plants were incubated with purified starch granules from Arabidopsis WT. Proteins bound to starch were eluted and LCI9-mCherry was detected in the starch bound fraction using immunoblotting (Fig. **S12**). To assess for changes to starch granule morphology, isolated starch was imaged using scanning electron microscopy. Starch granules from both the WT and the LCI9-mCherry expressing plants were lenticular although starch from the LCI9-mCherry was slightly smaller on average (Fig. **4c,d**). LCI9 therefore interacts with plant starch but does not dramatically alter its morphology.

**Figure 4.**
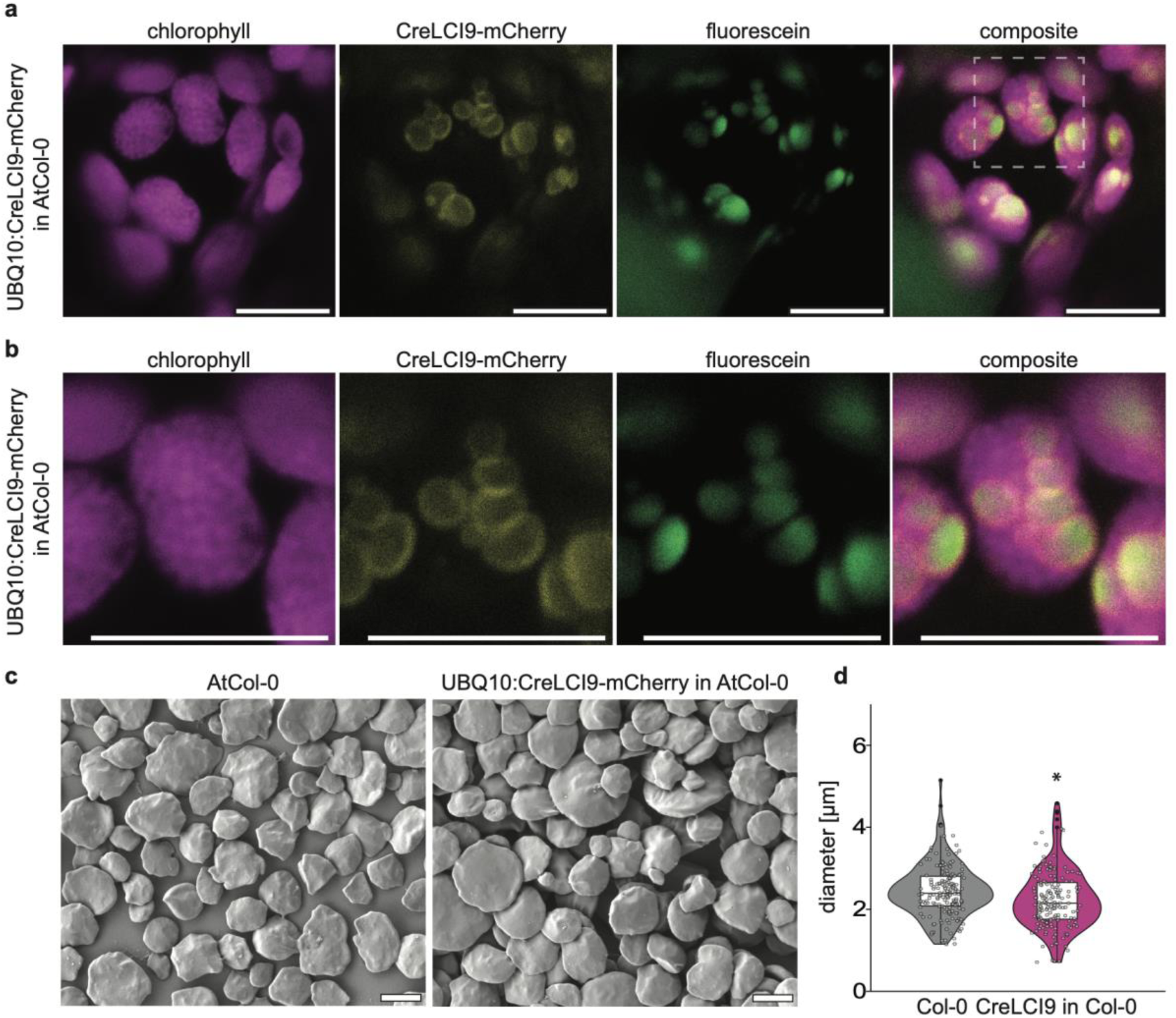
CreLCI9 interacts with starch granules and decreases starch granule size in planta. **(a-b)** Confocal laser scanning images of Arabidopsis (At) mesophyll cell chloroplasts of 21 day old plants, in a line stably overexpressing CreLCI9-mCherry (line 1) shown in yellow. Starch granules were stained with fluorescein shown in green. Chlorophyll fluorescence is shown in magenta. Arrows indicate example starch granules.Dashed square indicates enlarged region in (b). Bars: 10 μm. **(c)** Scanning Electron Micrographs of purified starch granules. Bars: 2 μm. **(d)** Size distribution of purified starch granules from a pool of 60 28 day old rosettes. Diameter was measured in ImageJ (200 particles were measured). Significant differences under Mann-Whitney Rank Sum Test are indicated with an asterisk (*) (P**≦** 0.001). For all boxplots, the bottom and top of the box represent the lower and upper quartiles, respectively, and the band inside the box represents the median. The ends of the whiskers represent values within 1.5x of the interquartile range, whereas values outside are outliers.

## Discussion

Appropriate assembly of a starch sheath that encapsulates the pyrenoid is critical for an efficient pCCM in Chlamydomonas (Toyokawa *et al*., 2020). The starch sheath is a dynamic structure and is fully formed under CO_2_-limiting conditions and degraded under CO_2_-replete conditions (Ramazanov *et al*., 1994). During starch sheath formation, the total amount of starch does not change suggesting that stromal starch granules are relocalised to the pyrenoid and remodelled and/or are degraded as pyrenoid starch plates are formed *de novo* (Izumo *et al*., 2011). In addition to their concave shape, the starch sheath plates also need to have gaps to enable membrane connections between the thylakoid network and the pyrenoid tubules. Thus, pyrenoid starch sheath formation likely requires careful regulation.

Here we have shown that LCI9 is a key component necessary for correct formation of the starch sheath in Chlamydomonas. In cells where the *LCI9* gene is disrupted, the starch plates had a non-uniform shape often with pits or pores, a phenomenon associated with increased susceptibility to degradation (Benmoussa *et al*., 2006). Although our extraction was performed at 4 ℃, it is not clear if this degradation is occurring *in vivo* or post extraction. This increased susceptibility to degradation could be due to altered starch composition, the increased surface area or the composition of proteins that coat the starch. The starch also featured protruding growths, which could be associated with high-amylose starch (Ahmed *et al*., 2012). By generating a proxiome for LCI9 using TurboID we found that LCI9 is spatially close to both starch synthesis and starch degradation enzymes (Fig. **3c**, Table **S4,S5**). It should be noted that this proxiome may simply represent the local starch sheath proteome and does not necessarily represent the interactome of LCI9. However, in combination with SBE3 being found to interact with LCI9 by Mackinder *et al*. (2017), our data supports that LCI9 likely interacts with or influences the regulation of both starch synthesis and degradation.

In plants, a group of proteins known as PTST were identified as essential for guiding starch granule initiation (Seung *et al*., 2015; 2017). LCI9 has a similar domain structure to PTST proteins by containing both a CBM and a coiled-coil domain and therefore might serve a similar scaffolding function. However, PTST proteins have CBM48 domains rather than the CBM20 domains found in LCI9 and the other genes in the LCI9 gene cluster. LCI9 forms a distinct phylogenetic group with genes from this cluster suggesting that these proteins have a distinct role to PTSTs. Our generated proxiome suggests that LCI9 is closely associated with multiple starch remodelling enzymes (Fig. **3**), which, in combination with the localisation of LCI9, suggests the function of LCI9 may be recruiting or regulating starch biosynthesis enzymes at the starch sheath specifically.

Starch morphology can be influenced by physical aspects such as the thylakoid membrane structure (Esch *et al*., 2022; Isle *et al*., 2025). The spherical Rubisco matrix may provide a physical guide to starch plate formation, enabling the synthesis of concave plates on the surface of the matrix. This is supported by LCI9 appearing to bind to the surface of starch granules facing the Rubisco matrix during the first hour of formation in several of the imaged pyrenoids (Fig. **2f****; S8**), and the physical proximity of LCI9 to SAGA2, which localises at the starch-Rubisco matrix interface (Fig. **3c**, Meyer *et al*., 2020). In cells lacking the ability to form the phase separated Rubisco matrix, starch granules are still abundant at the canonical pyrenoid site but are round and not shaped into plates, suggesting that the pyrenoid matrix is required for the shaping of pyrenoid starch (Mackinder *et al*., 2016; He *et al*., 2020).

Pyrenoid starch plates and the stromal starch granules appear to be under separate control and LCI9 may be part of a broader class of proteins that specifically control pyrenoid starch sheath formation (Ramazanov *et al*., 1994; Findinier *et al*., 2019). LCI9 is associated with two proteins that have already been implicated in pyrenoid starch specifically, BSG1 (Cre02.g091750; Findinier *et al*., 2019) and Cre03.g187150 (Hennacy, 2023). Cells deficient in BSG1 over accumulate pyrenoid starch rather than switch to stromal starch production under nitrogen starvation, suggesting that BSG1 is required for transitioning between the two types of starch (Findinier *et al*., 2019). The Cre03.g187150 gene has not been extensively studied in the context of starch but it was observed that when knocked out alongside *SAGA1*, overly large starch granules accumulate adjacent to Rubisco condensates (Hennacy, 2023).

From our time course images, after a transition into low CO_2_, LCI9 appears to first localise at the interface between the pyrenoid associated starch granules and the pyrenoid matrix. SAGA2 also localises at this interface (Meyer *et al*., 2020), and through interacting with LCI9 could initially recruit LCI9 to this interface. However, LCI9 subsequently localises all the way around pyrenoid starch before finally becoming enriched in the gaps between different starch plates (Fig. **2g,h**) (Mackinder *et al*., 2017). LCI9 may guide the lateral growth of starch as the granules become more plate-like. Once two plates meet, LCI9 could form part of a stabilizing complex to prevent further growth or degradation. Without LCI9 protecting the starch sheath, starch synthesis and degradation enzymes may gain unregulated access to the starch plates resulting in the aberrant protrusions and pits we observed via SEM (Fig. **2e**). Collectively our data supports that LCI9 plays a critical role in starch sheath formation, potentially controlling the recruitment of starch metabolism enzymes to pyrenoid starch and stabilising plate-plate interfaces.

Engineering a starch sheath around a reconstituted proto-pyrenoid in plants is a key next step towards building a functional pCCM (Atkinson *et al*., 2020; Adler *et al*., 2022; Fei *et al*., 2022). While SAGA1 and SAGA2 appear sufficient for recruiting starch (Atkinson *et al*., 2024), other proteins are likely needed to shape the starch into plates. Our preliminary work showed that LCI9 interacts with chloroplastic starch granules in Arabidopsis but that the starch granules remained morphologically similar to WT plants (Fig. **4c**). Starch granules from LCI9 expressing plants tended to be slightly smaller, perhaps due to LCI9 blocking access to starch synthesis enzymes by steric hindrance (Fig. **4d**). The lack of major morphological differences could be due to the absence of LCI9 interaction partners that are not present in Arabidopsis. Investigating the impact of expressing LCI9 in Arabidopsis lines with proto-pyrenoids could be an exciting next step towards engineering a pyrenoid starch sheath in plants.

## Acknowledgements

The authors thank Imaging and Cytometry Technology Facility at the University of York for imaging support. Specifically, Karen Hodgkinson for SEM, Clare Steele-King for TEM and Grant Calder for light microscopy support. Stephen R. Mitchell from the University of Edinburgh also supported TEM work. Adam Dowle from the University of York Centre of Excellence in Mass Spectrometry for supporting proximity labelling-mass spectrometry experiments. The work was funded by a UKRI-Future Leaders Fellowship (MR/T020679/1 and MR/Y034074/1) and EPSRC (EP/W024063/1), BBSRC (BB/R001014/1), BBSRC IAA grants to LM; and a Bill and Melinda Gates Foundation and FCDO (INV-054558), Carbon Technology Research Foundation (AP23-1_023), and BBSRC-NSF/BIO (BB/Y000323/1, BB/S015337/1) grants to LM and AMc.

## Data Availability

The majority of data that supports the findings of this study are available in the supplementary material of this article. Raw mass spectrometry data will be deposited at MassIVE. Strains and plasmids used in this will be deposited with the Chlamydomonas Resource Center. Any additional data will be made available upon request.

## Author Contributions

LA, CSL, LE, AMc and LM conceived the study. JB and GY did initial validation of mutants and generated plasmids. LA generated and validated complemented lines, performed microscopy for Chlamydomonas and isolated starch for SEM. LA and CSL performed turbo ID, CSL analysed the data. CSL performed TEM and immunoblots. LE performed all plant based work and phylogenetic analysis. OD performed solid media growth assays and O_2_ electrode measurements. SG assisted with microscopy. LA wrote the manuscript with input from all authors.

## Supplemental Data

**Supplemental Figure S1.**
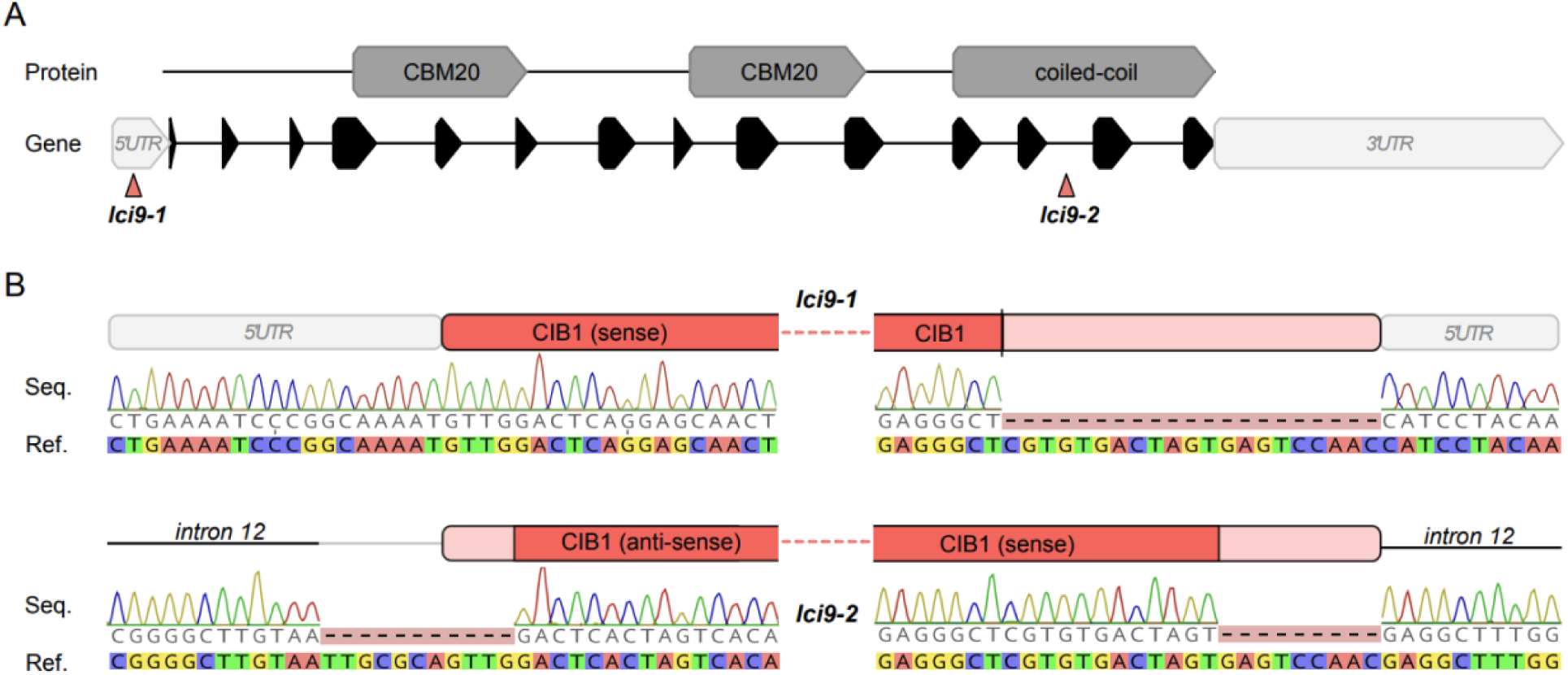
Validation of CIB1 cassette insertion. **(a).** Schematic of cassette insertion to LCI9 gene. **(b).** Sequencing data of CIB1 cassette junctions in *lci9-1* and *lci9-2*. Dark red indicates verified sequence of the inserted CIB1 cassette, pale red shows where CIB1 cassette has been truncated. For *lci9-2*, a genomic deletion in intron 12 upstream of the CIB1 cassette is shown by the thin grey line.

**Supplemental Figure S2.**
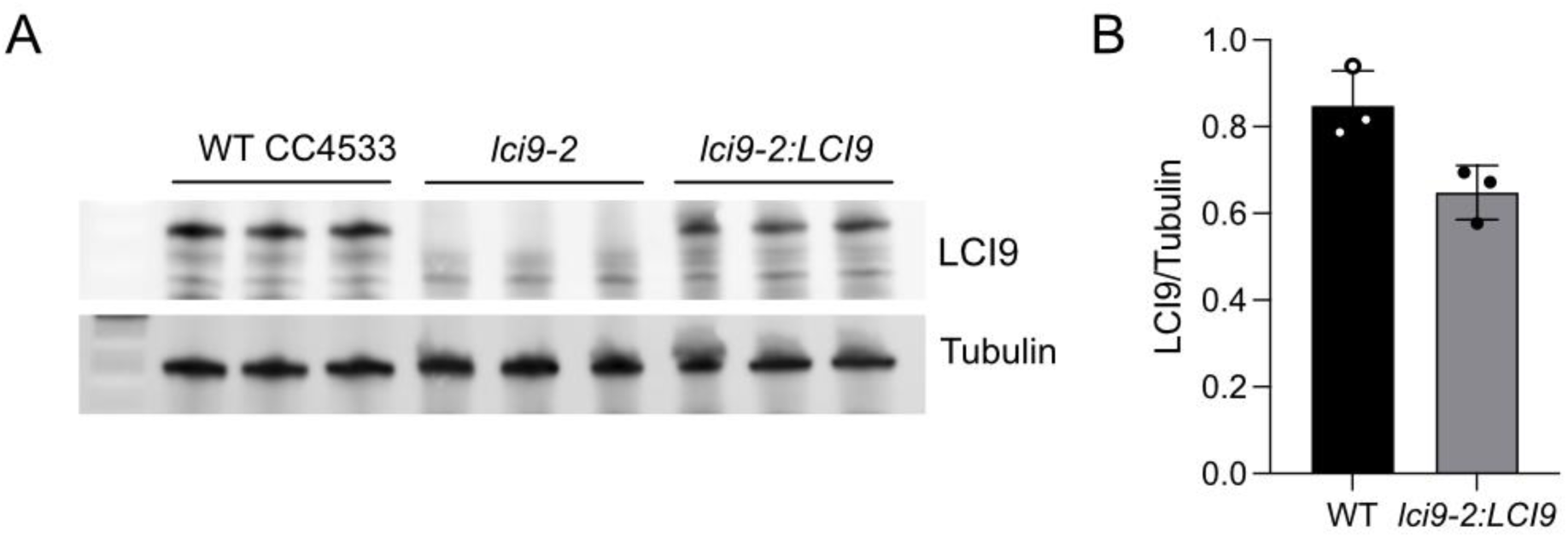
LCI9 levels in wild-type compared to lci9-2:LCI9 complemented line. **(a).** Immunoblot of LCI9 and tubulin in three biological replicates. **(b).** Relative abundance of LCI9 protein in wild-type compared to *lci9-2:LCI9*.

**Supplemental Figure S3.**
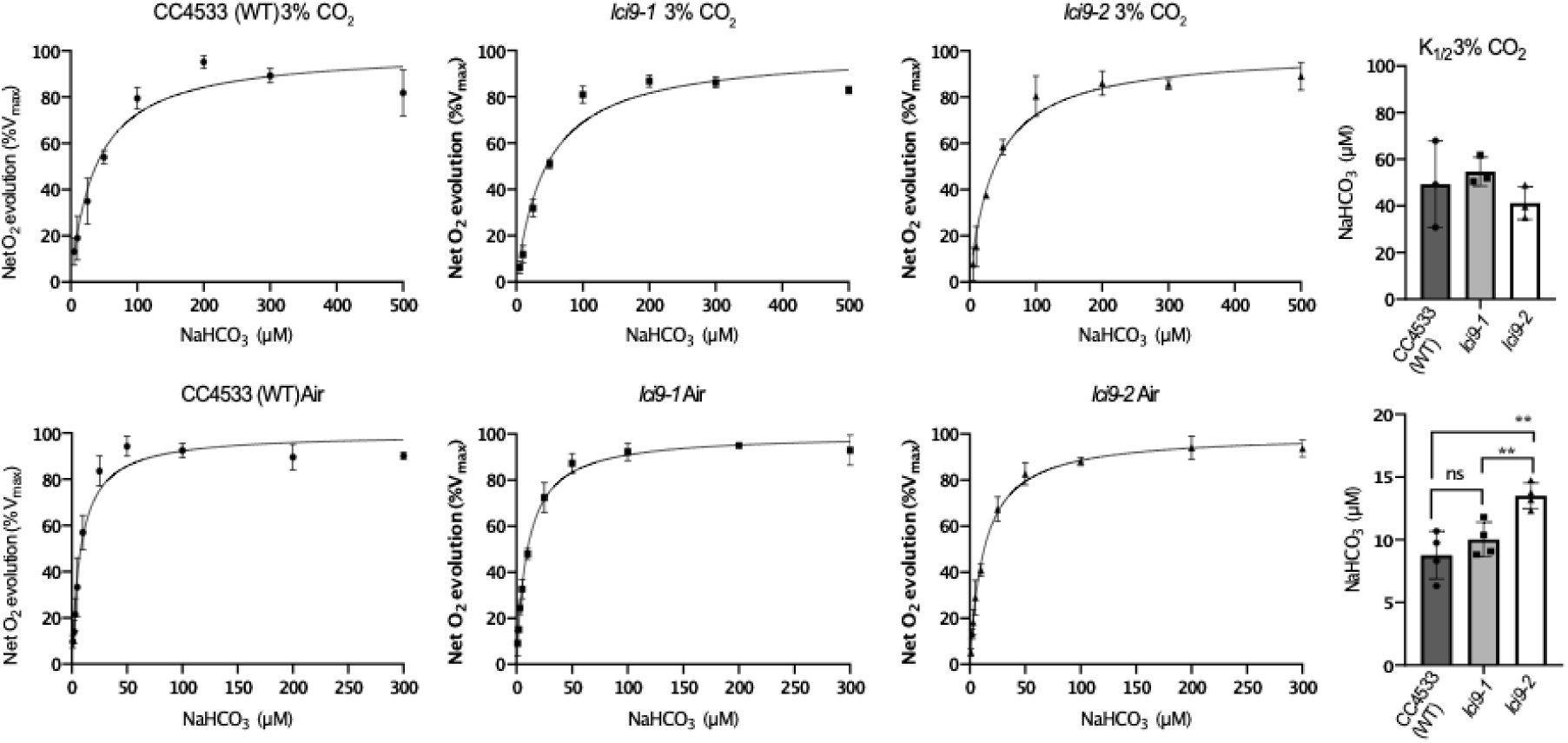
Ci affinity of *lci9-1* and *lci9-2* compared to control strain cc-4533.

**Supplemental Figure S4.**
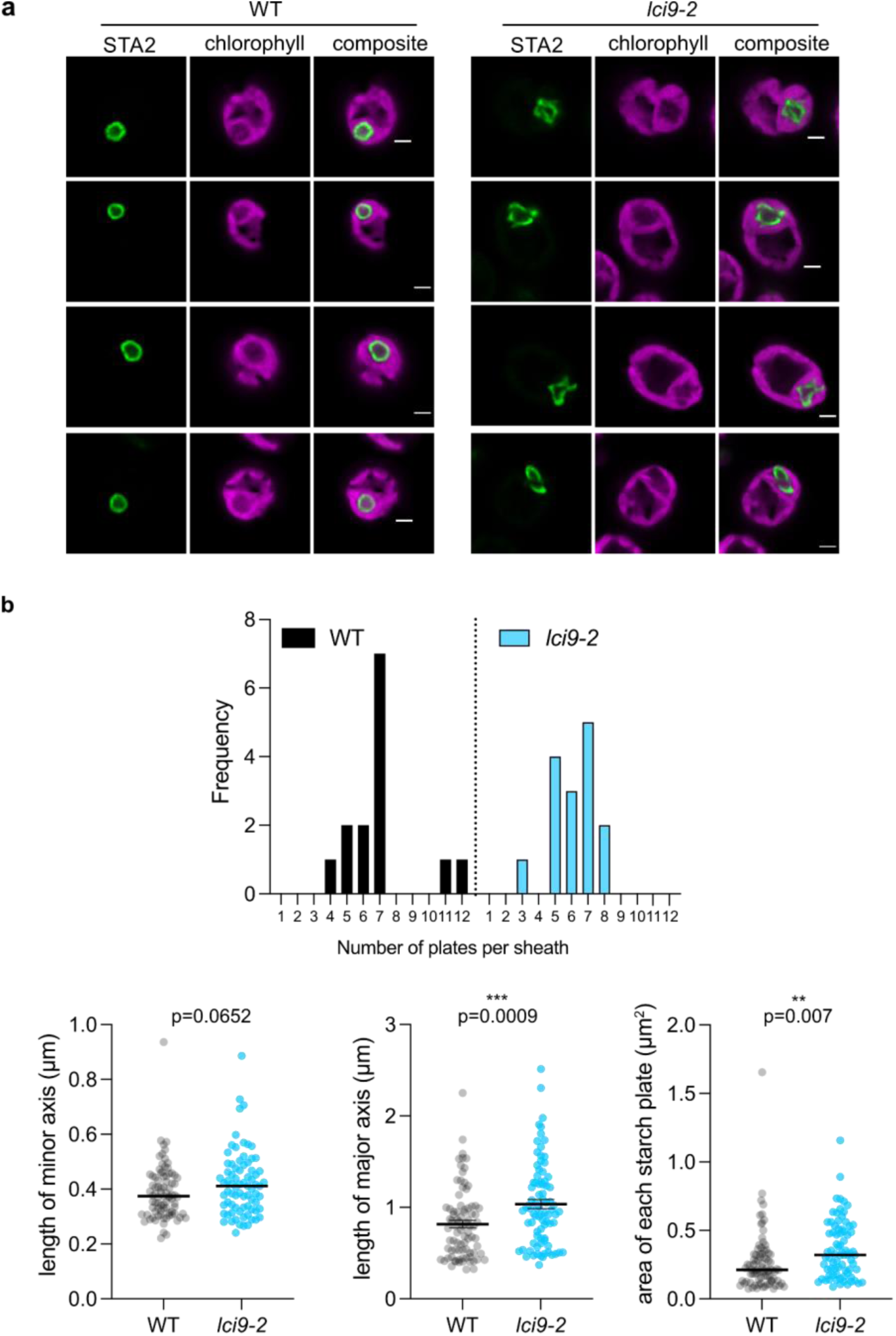
Extended data for Fig. 2. **(a)** Additional images of STA2-mCherry expressed in WT and *lci9-2.* Cells are adapted to low (0.04%) CO_2_. Images are single z planes through the middle of the pyrenoid. Scale bar is 2 µm. **(b)** Characteristics of individual starch plates in WT and *lci9-2* across 25 different cells. Asterisks represent significant differences as determined by an unpaired, two-tailed t-test.

**Supplemental Figure S5.**
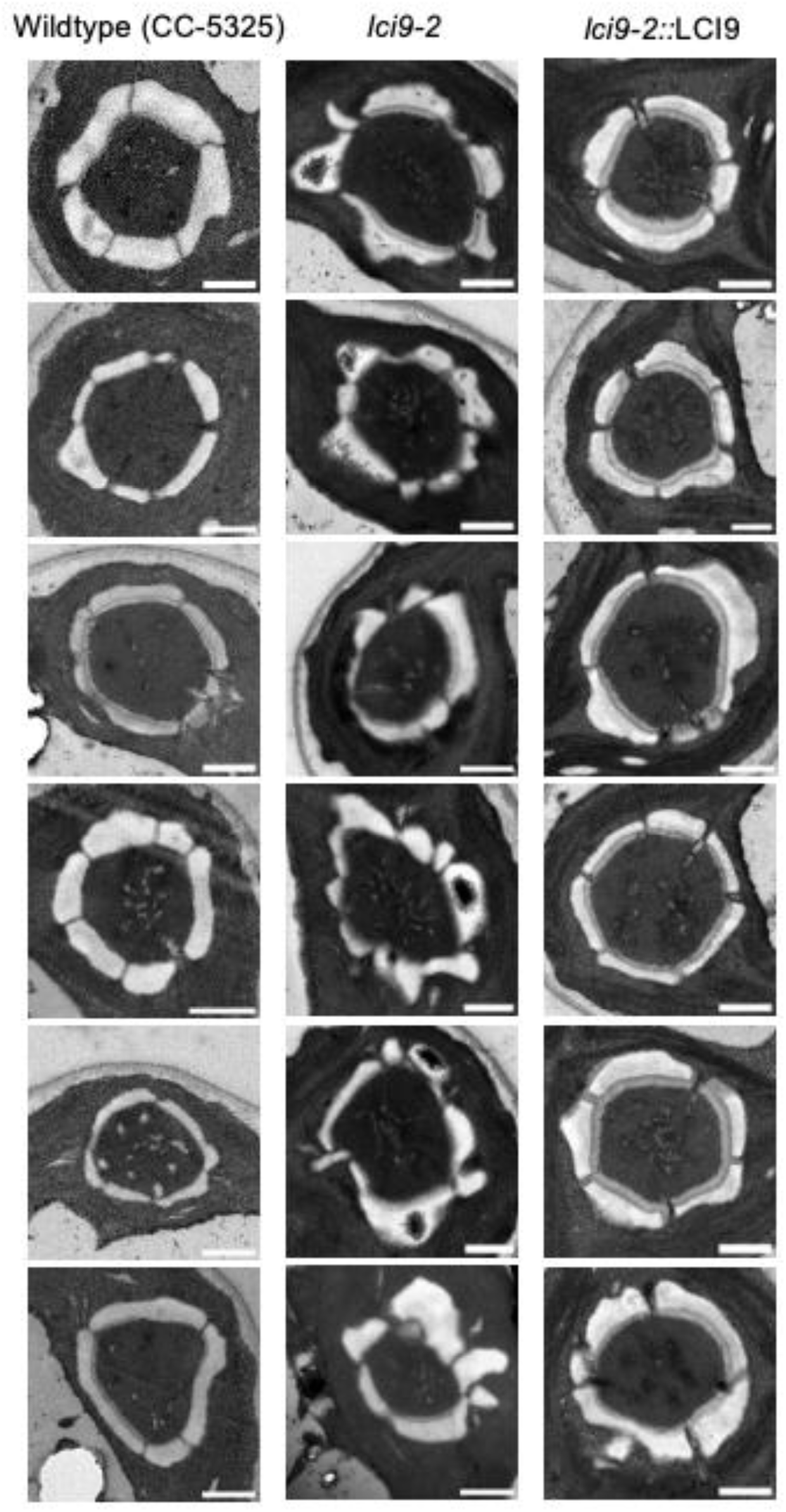
Additional transmission electron micrographs in support of Figure 3D. Cells were harvested under low (0.04%) CO_2._ Scale bar is 1 µm.

**Supplemental Figure S6.**
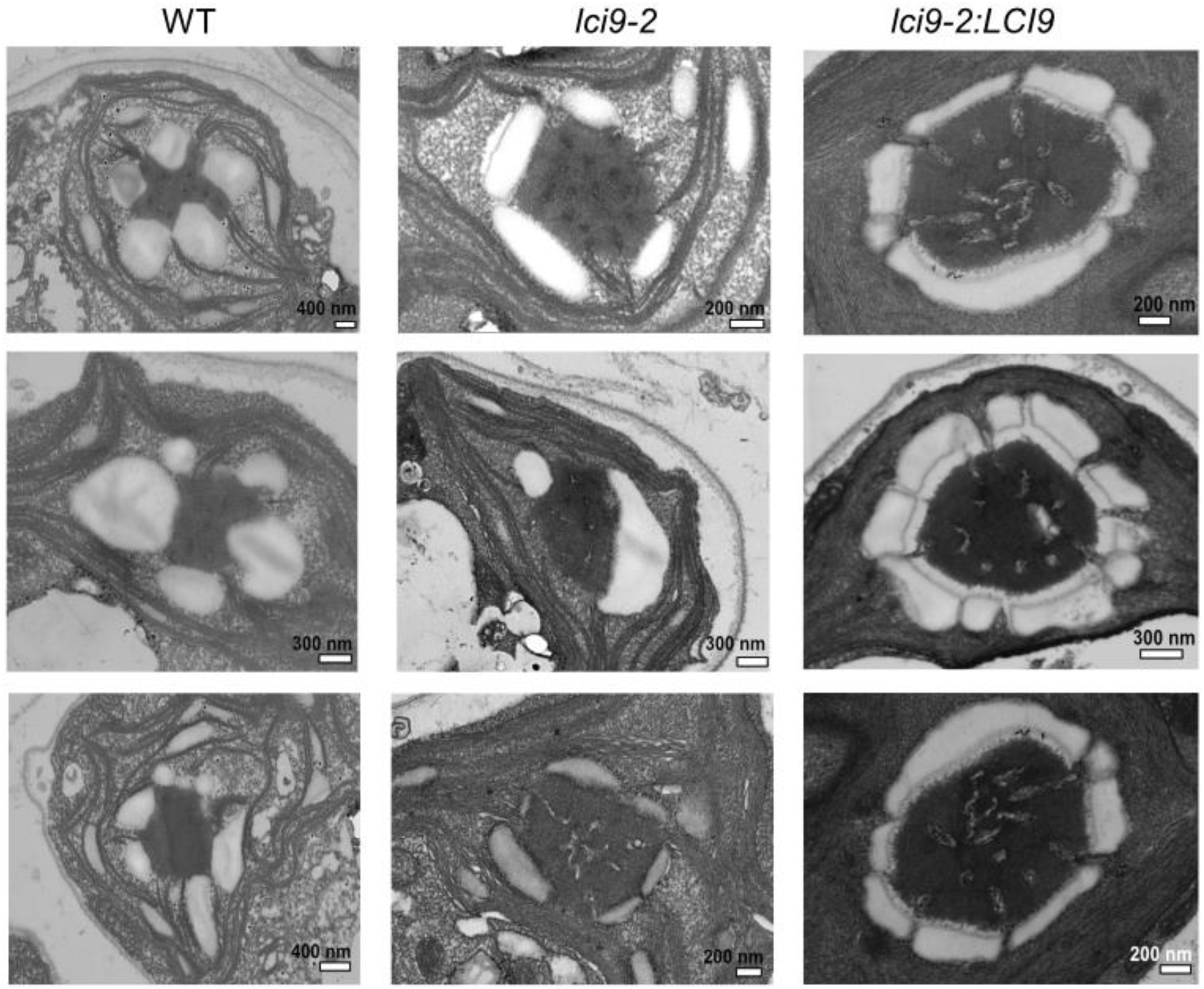
TEM of pyrenoids of cells at high CO_2_. Cells were harvested under high (3%) CO_2._ Scale bars are indicated in each image.

**Supplemental Figure S7.**
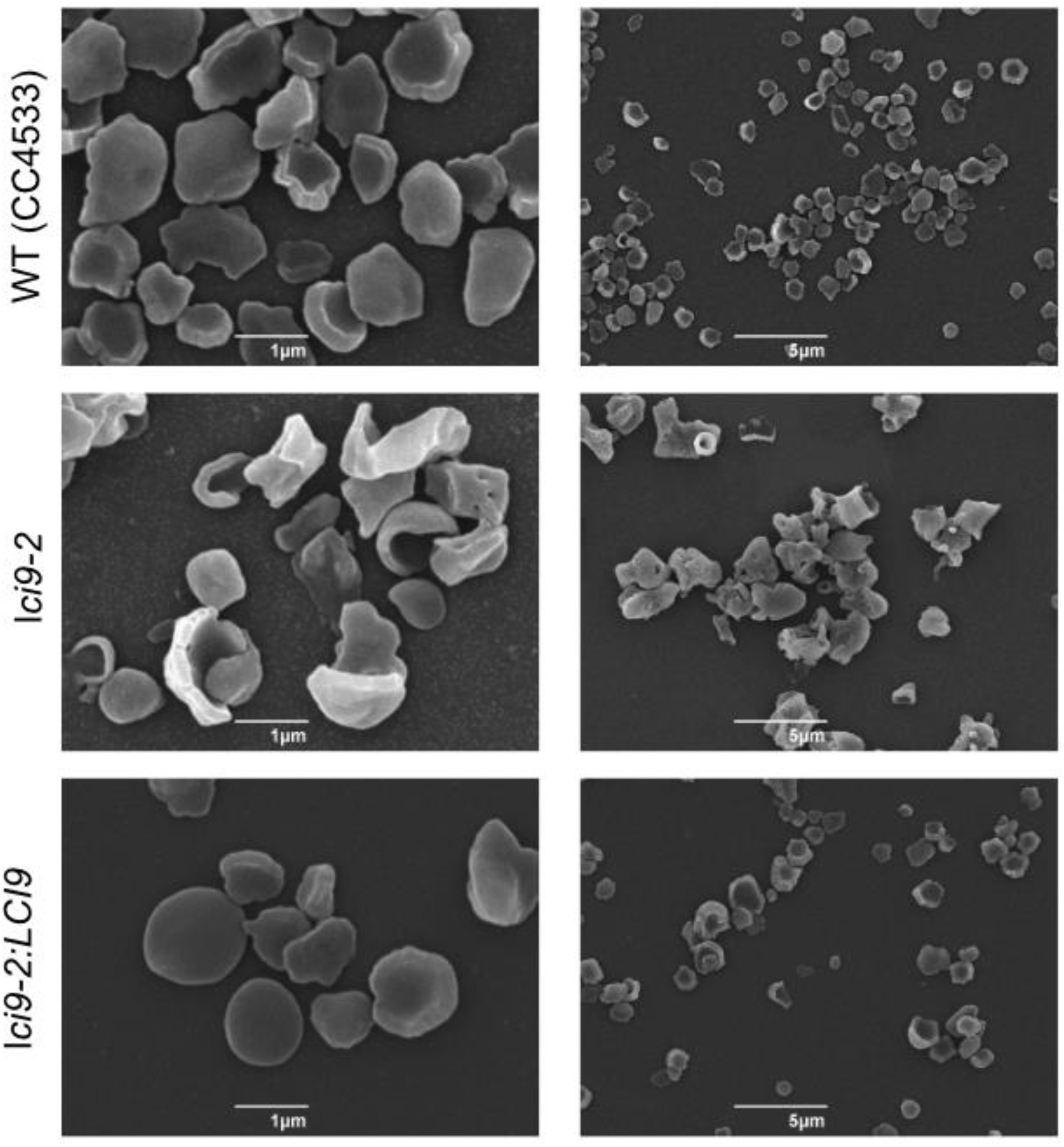
Additional SEM images of starch isolated from WT, *lci9-2* and *lci9-2:LCI9* cells grown under low (0.04%) CO_2_ conditions. Supports figure 2F.

**Supplemental Figure S8.**
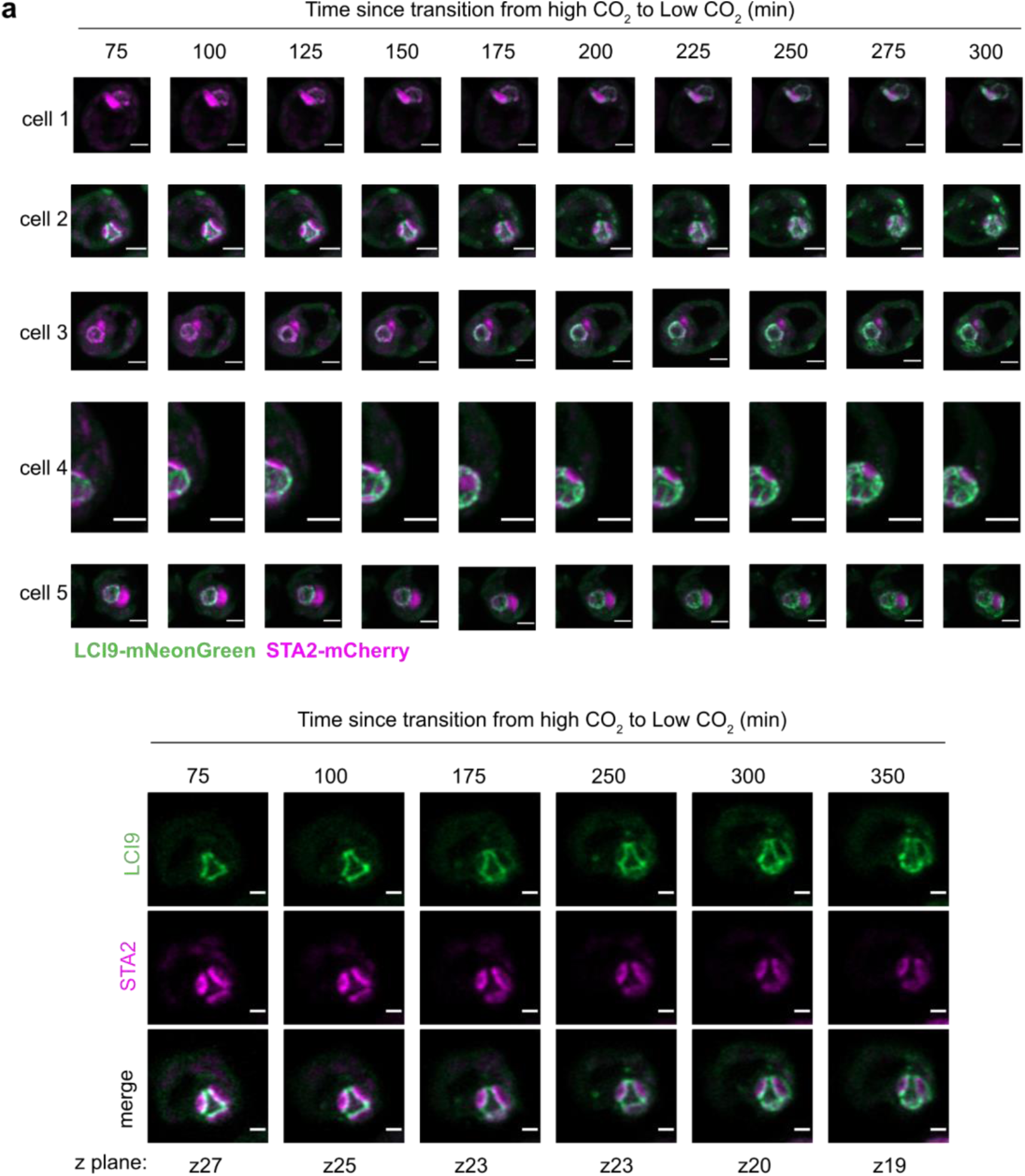
Additional cells from timelapse imaging of high CO_2_ to low CO_2_ transition in support of Figure 2G. **(a)** Maximum z projections of cells with LCI9-mNeonGreen and STA2-mCherry shown in green and magenta, respectively. Scale bars are 1 µm. **(b)** A single z plane of cell 2. The z plane has been adjusted to keep the cell in focus to compensate for drift during the experiment. Scale bar is 1 µm.

**Supplemental Figure S9.**
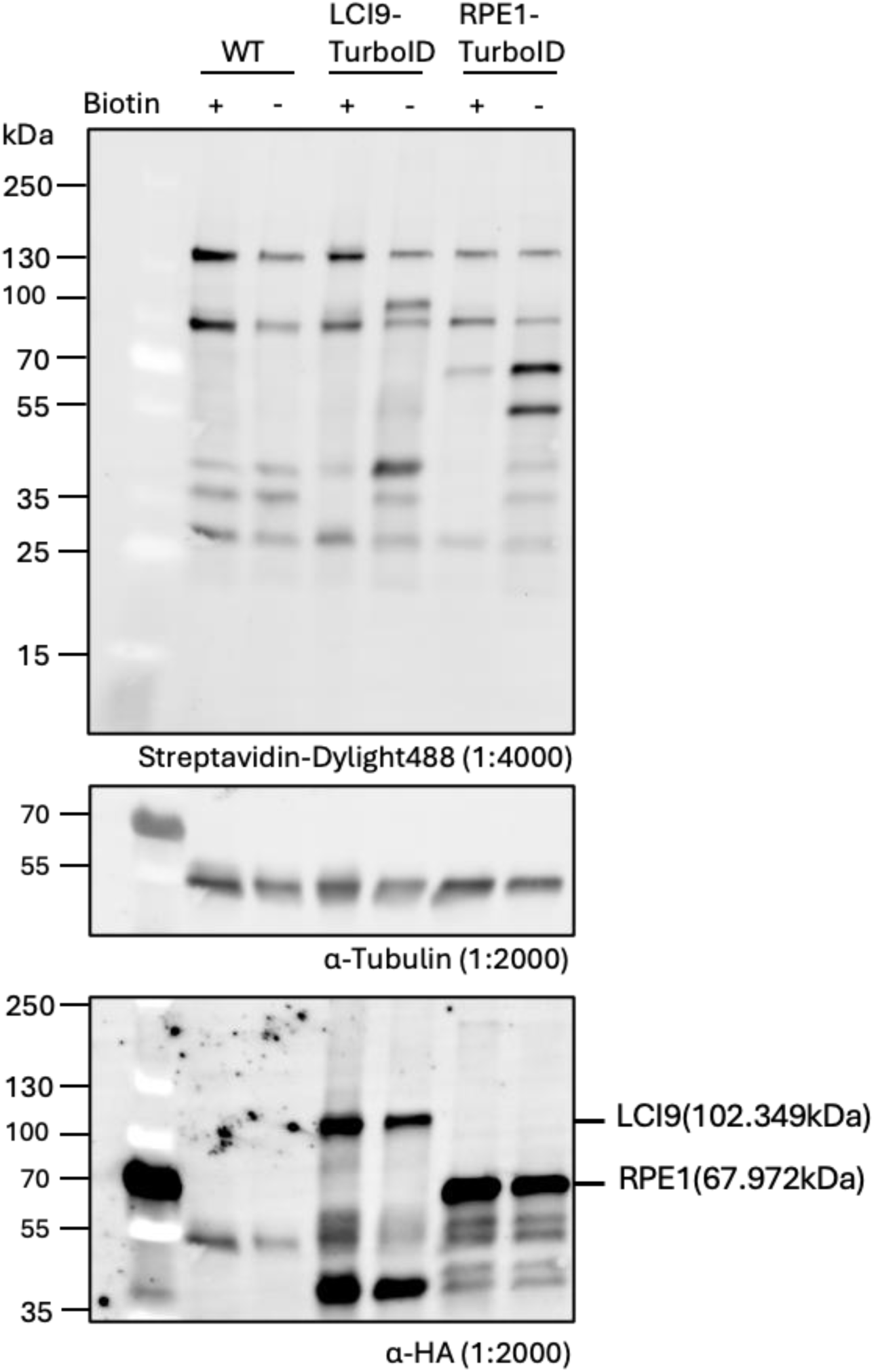
Confirmation of expression and activity of TurboID constructs. The biotinylation activity of the TurboID expression strains, as determined in the absence (−) or presence (+) of 2.5 mM biotin for 2 h. Cultures are separately harvested both before and after the biotin labelling treatment. Biotinylation was visualized via immunoblotting whole-cell lysate with a streptavidin conjugate. The abundance of LCI9-TurboID (102 kDa) and RPE1-TurboID (67kDa), was probed by α-HA. α-tubulin was used as a loading control.

**Supplemental Figure S10.**
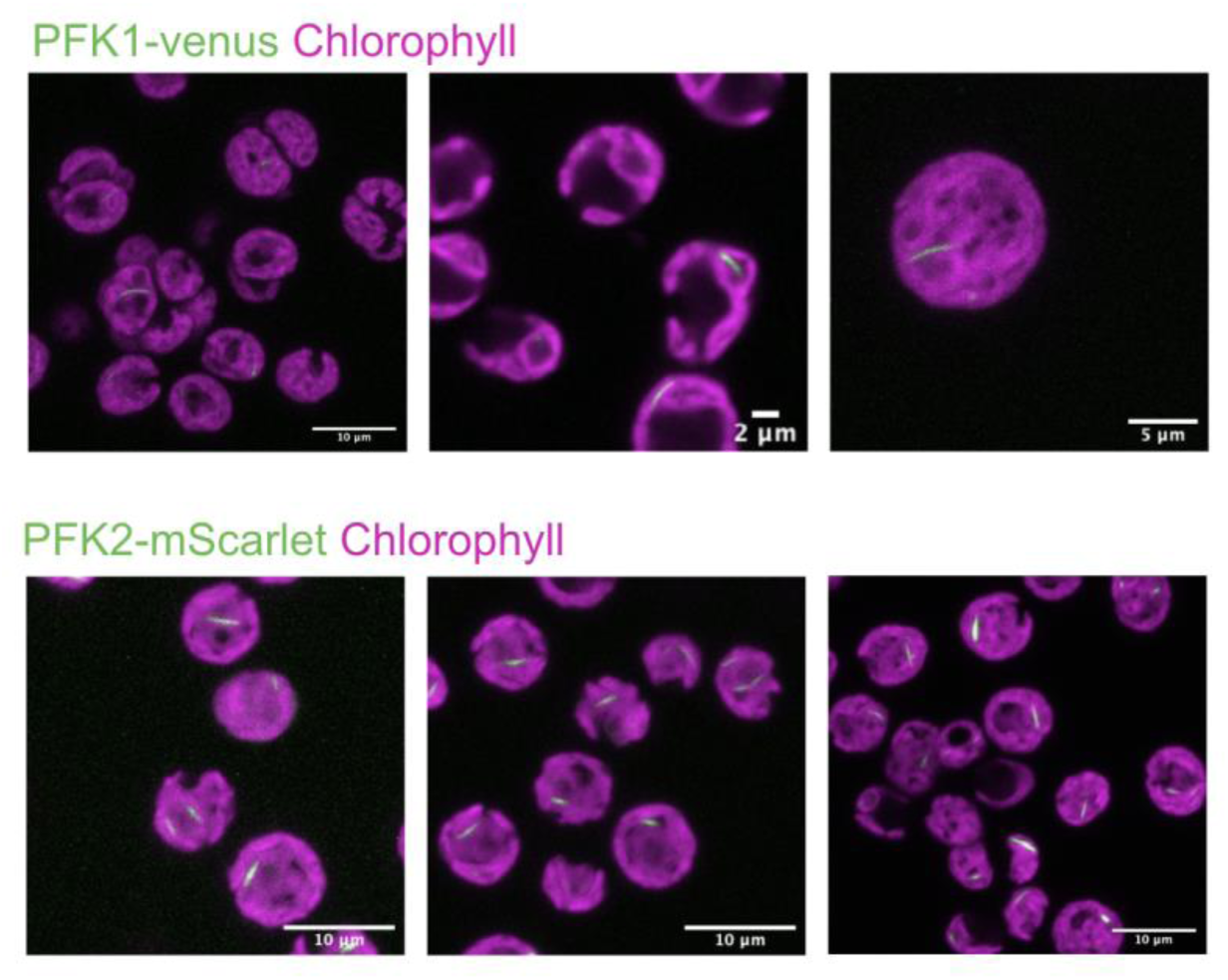
Localisation of PFK1 and PFK2 via fluorescent tagging.

**Supplemental Figure S11.**
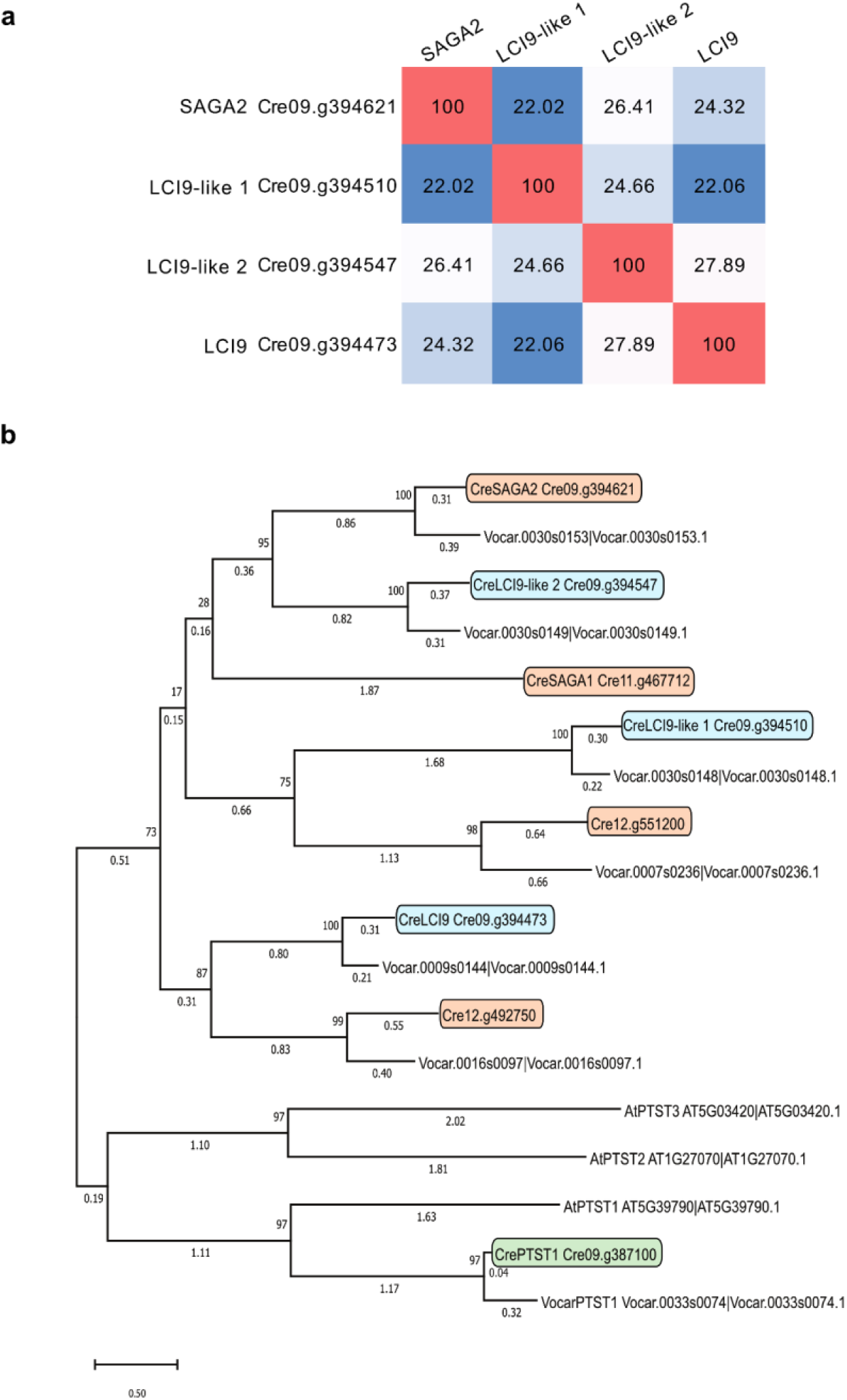
LCI9-like gene cluster phylogenetic analysis. LCI9 is part of a gene cluster including two other LCI9 homologues. **(a)** Percentage identity matrix of amino acid sequences of the LCI9 gene cluster. **(b)** Phylogenetic tree of CBM-coiled coil proteins across selected green algae and Arabidopsis. LCI9 like genes, other CBM20 proteins and PTST are shown in blue, orange and green, respectively. The tree with the highest log likelihood (-45,002.51) is shown. The percentage of trees in which the associated taxa clustered together is shown next to the branches. Initial tree(s) for the heuristic search were obtained automatically by applying Neighbor-Join and BioNJ algorithms to a matrix of pairwise distances estimated using the JTT model, and then selecting the topology with superior log likelihood value. The tree is drawn to scale, with branch lengths measured in the number of substitutions per site (next to the branches).

**Supplemental Figure S12.**
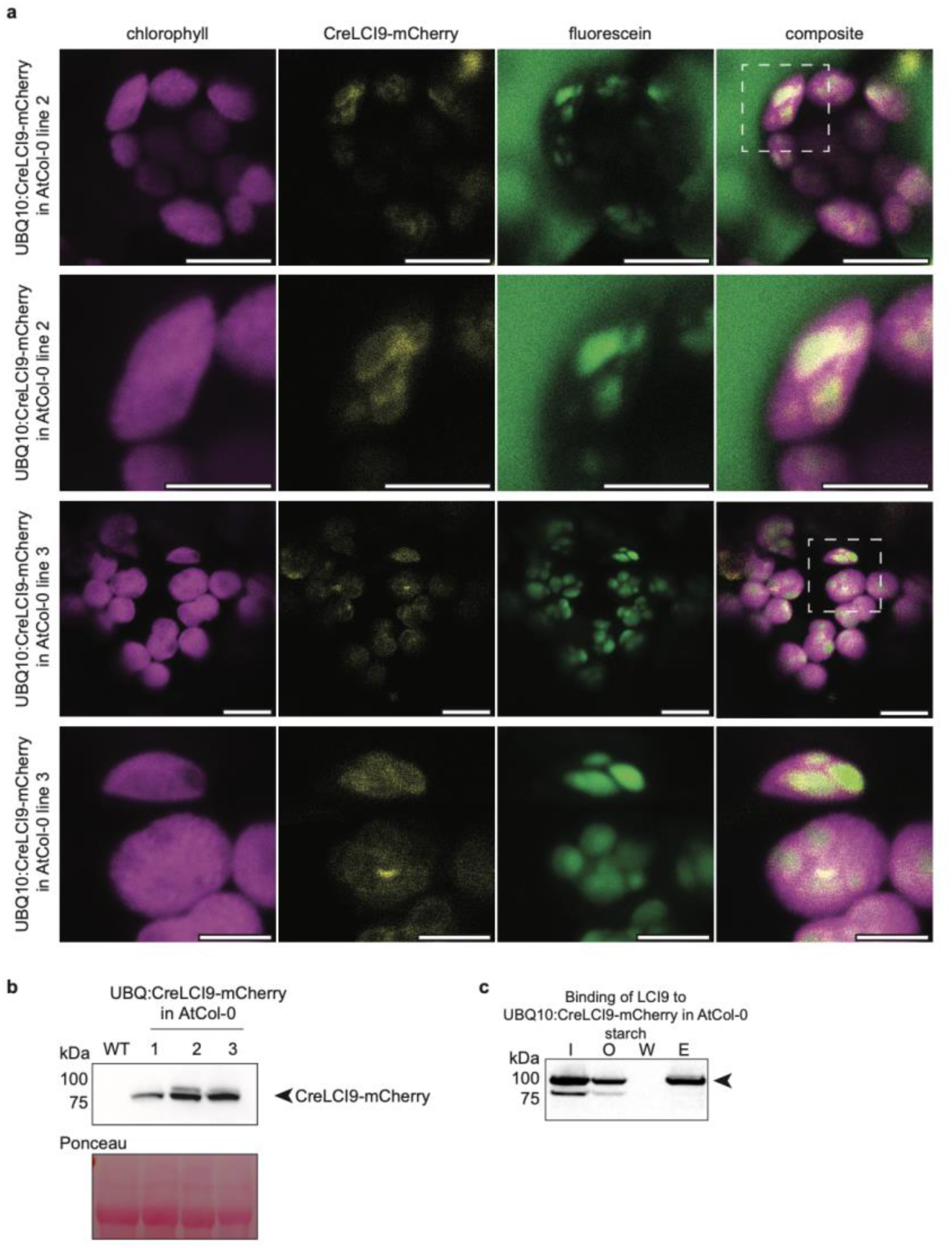
CreLCI9 is expressed in 3 independent Arabidopsis lines and interacts with starch granules in planta. **(a)** Confocal laser scanning images of Arabidopsis (At) mesophyll cell chloroplasts of 28 day old plants, in independent lines stably overexpressing CreLCI9-mCherry shown in yellow. Starch granules were stained with fluorescein shown in green. Chlorophyll fluorescence is shown in magenta. Dashed squares indicate enhanced regions below. Bars: 10 µm and 5 µm for close ups. **(b)** Immunoblot using mCherry antibody to detect CreLCI9-expression in 2-3 representative plants of AtCol-0 controls and UBQ:CreLCI9-mCherry in AtCol-0 lines 1,2 and 3. **(c)** Starch binding assay for CreLCI9-mCherry using purified starch from UBQ10:CreLCI9-mCherry in AtCol-0 line1 and protein extracts from 28 day old UBQ10:CreLCI9-mCherry in AtCol-0 line 1 leaves. Input (I) indicates total protein extract. Output (O) indicates supernatant after incubation with starch granules. Wash (W) indicates supernatant from the last wash step and Elute (E) indicates supernatant after protein elution from starch granules. Arrows indicate CreLCI9-mCherry. CreLCI9-mCherry was detected using a mCherry-antibody (ab213511, abcam).

**Supplemental Table 1.** Plasmids used in this study.

| Name | Organism |
| --- | --- |
| STA2-mCherry | Chlamydomonas |
| LCI9-mNeonGreen | Chlamydomonas |
| LCI9 | Chlamydomonas |
| LCI9-Turbo | Chlamydomonas |
| RPE1-Turbo | Chlamydomonas |
| LCI9-mNeonGreen/STA2-mCherry | Chlamydomonas |
| LCI9-mCherry, KanR | Arabidopsis |

**Supplemental Table 2.**
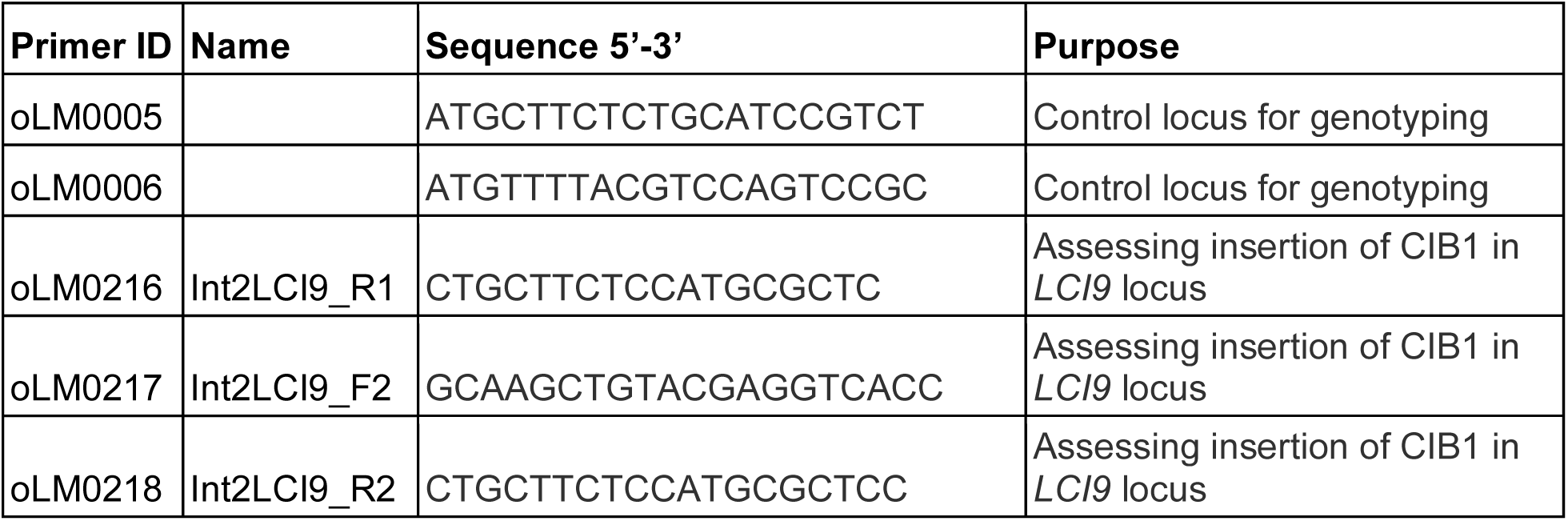
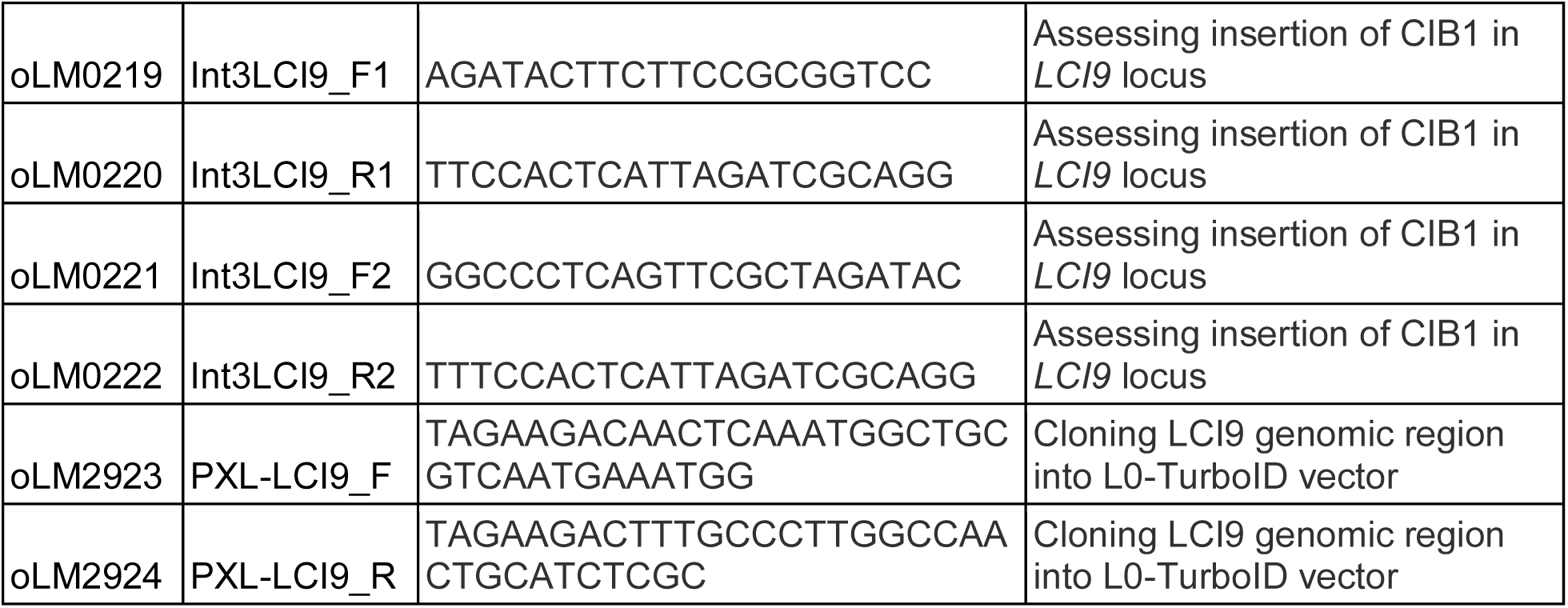
Primers used in this study.

**Supplemental Table 3.** Strains used in this study.

| Strain | Background | Reference |
| --- | --- | --- |
| *lci9-1* LMJ.RY0402.123483 | CC-4533 | Li *et al.* (2019) |
| *lci9-2* LMJ.RY0402.172898 | CC-4533 | Li *et al.* (2019) |
| *lci9-2:LCI9* (E4) | *lci9-2* | This study |
| WT<sub>cc-4533</sub>:STA2-mScarlet-I | CC-4533 | This study |
| *lci9-2*:STA2-mScarlet-I | *lci9-2* | This study |
| WT<sub>cc-4533</sub>:*LCI9-mNeonGreen/STA2-mCherry* | CC-4533 | This study |
| *epyc1* |  | Mackinder *et al.* (2016) |
| WT<sub>cc-4533</sub>:LCI9-Turbo | CC-4533 | This study |
| WT<sub>cc-4533</sub>:RPE1-Turbo | CC-4533 | Lau *et al.* (2023) |

**Supplemental Table 4.**
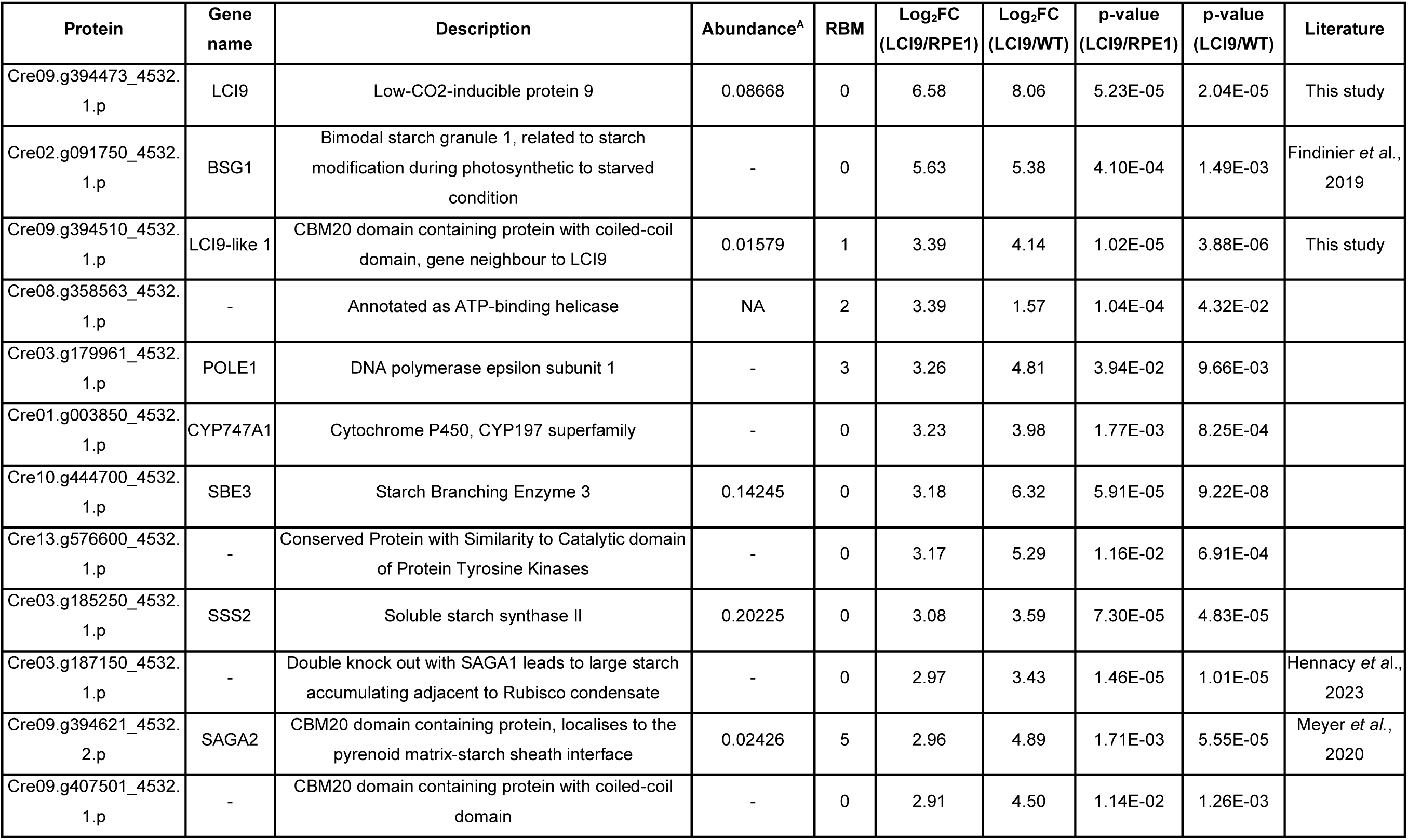
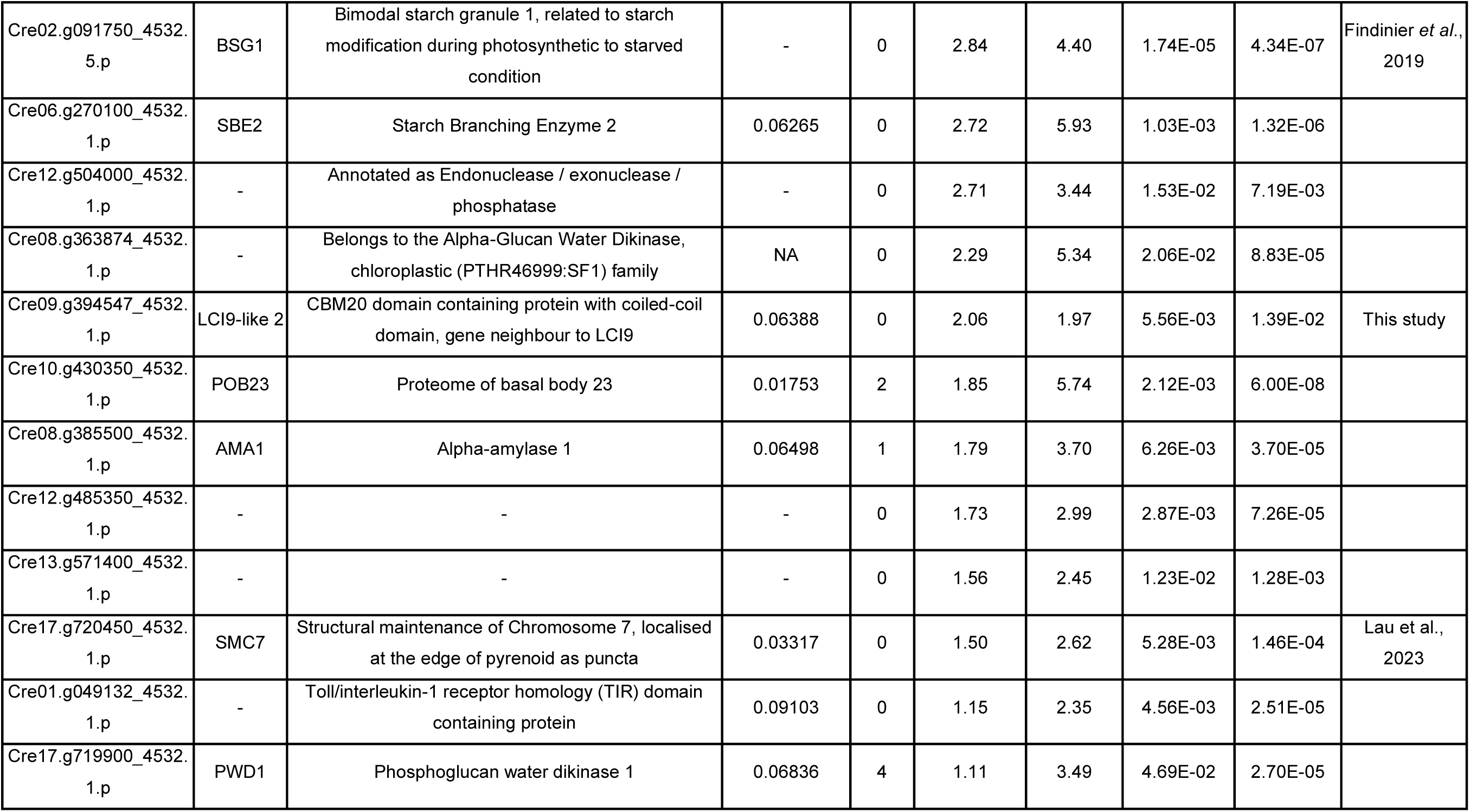
Identified proteins in LCI9 high-confidence proxiome. The LCI9-TurboID preferentially labelled proteins found to be above the enrichment threshold of a Log_2_FC value >1 and p-value <0.05 when comparing against both the RPE1-TurboID and Wildtype control. The protein abundance listed refers to the determined protein abundance of cc-1690 grown under the control condition obtained from Arend *et al*., 2023.

